# Transient DNA Binding Induces RNA Polymerase II Compartmentalization During Herpesviral Infection Distinct From Phase Separation

**DOI:** 10.1101/375071

**Authors:** David T McSwiggen, Anders S Hansen, Hervé Marie-Nelly, Sheila Teves, Alec B Heckert, Claire Dugast-Darzacq, Yvonne Hao, Kayla K Umemoto, Robert Tjian, Xavier Darzacq

**Affiliations:** Department of Molecular and Cell Biology, University of California Berkeley, CA, USA, 94720; California Institute of Regenerative Medicine Center of Excellence, University of California Berkeley, CA, USA, 94720; Department of Biochemistry and Molecular Biology, University of British Columbia, Vancouver, BC, Canada, BC V6T 1Z4; Howard Hughes Medical Institute, University of California Berkeley, CA, USA, 94720

**Keywords:** Transcription, Phase Separation, Nuclear Organization, Single Particle Tracking, RNA Polymerase II, Herpesvirus, Microscopy

## Abstract

During lytic infection, Herpes Simplex Virus 1 generates replication compartments (RCs) in host nuclei that efficiently recruit protein factors, including host RNA Polymerase II (Pol II). Pol II and other cellular factors form hubs in uninfected cells that are proposed to phase separate via multivalent protein-protein interactions mediated by their intrinsically disordered regions. Using a battery of live cell microscopic techniques, we show that although RCs superficially exhibit many characteristics of phase separation, the recruitment of Pol II instead derives from nonspecific interactions with the viral DNA. We find that the viral genome remains nucleosome-free, profoundly affecting the way Pol II explores RCs by causing it to repetitively visit nearby binding sites, thereby creating local Pol II accumulations. This mechanism, distinct from phase separation, allows viral DNA to outcompete host DNA for cellular proteins. Our work provides new insights into the strategies used to create local molecular hubs in cells.

## Introduction

The ability to control the local concentration of molecules within cells is fundamental to living organisms. A classic example is the use of electrochemical gradients across membranes to facilitate cellular work. In recent years, our understanding of the forces driving the formation of sub-nuclear compartments has undergone a paradigm shift. A number of studies suggest that many proteins have the ability to spontaneously form separated liquid phases *in vitro* (Hyman et al., 2014), and recent work highlights the possibility that similar liquid compartments may occur *in vivo* (Shin et al., 2017; Strom et al., 2017). Such liquid-liquid demixing has been proposed to be a common mechanism to sequester specific macromolecules within a compartment, or to increase their local concentration to facilitate chemical interactions. Formation of these structures is thought to be predominantly driven by weak, multivalent protein:protein interactions mediated by intrinsically disordered regions (IDRs), which are often comprised of low complexity polypeptides, and sometimes aided by the presence of modular RNA or DNA binding motifs (Banani et al., 2017; Hyman et al., 2014).

These observations have opened new frontiers of inquiry in cell biology, and generated a deeper appreciation for the diversity of mechanisms that a cell may deploy to locally concentrate certain molecular components. The list of proteins—particularly nuclear proteins—that can undergo phase separation *in vitro* continues to grow (Courchaine et al., 2016). For example, recent studies of RNA Polymerase II (Pol II) and its regulators have shown that Pol II forms dynamic hubs whose size are dependent on the number of intrinsically disordered heptad peptide repeats contained within the C-terminal domain (CTD) (Boehning et al., 2018), and that various CTD interacting factors may form phase separated droplets in vitro (Lu et al., 2018). Even so, it remains unclear which molecular interactions drive the formation of domains via liquid-liquid phase separation (LLPS), or many different classes of interactions mediate such local molecular behavior (Chong et al., 2018). Importantly, we do not we fully understand the nature of the molecular forces that drive compartmentalization *in vivo*, still lack compelling evidence to establish the functional consequences of these compartments for the biological activities of their constituents.

Herpes Simplex Virus type 1 (HSV1) provides an attractive system to study this question because of its ability to form compartments in the nucleus of infected cells *de novo*. HSV1 is a common human pathogen that hijacks its host’s transcription machinery during lytic infection (Rice et al., 1994). Transcription of HSV1 genes occurs in three waves: immediate early, early, and late, with the latter strictly occurring only after the onset of viral DNA replication (Knipe and Cliffe, 2008). Like many DNA viruses, HSV1 creates subcellular structures called replication compartments (RC) where both viral and host factors congregate to direct replication of the viral genome, continue viral transcription, and assemble new virions (Schmid et al., 2014). Recent reports highlight the ability of HSV1 to usurp host Pol II to transcribe its own genome such that, once late gene transcription commences, the host chromatin is largely devoid of productively transcribing Pol II, and the majority of newly synthesized mRNAs are viral in origin (Abrisch et al., 2015; Rutkowski et al., 2015). Consistent with genomic data, immunofluorescence staining of infected cells shows a dramatic enrichment of Pol II and other nuclear factors in RCs (Rice et al., 1994).

Given this shift in both the sub-nuclear localization of Pol II upon infection, and its effect on the transcriptional output of an infected cell, we chose to examine the mechanism of Pol II recruitment to HSV1 RCs as a model case for the generation of new subcellular compartments. We employed a combination of imaging approaches, including live cell single particle tracking (SPT), fluorescence loss in photobleaching (FLIP), and fluorescence recovery after photobleaching (FRAP), in addition to single molecule localization microscopy to probe Pol II localization and behavior within RCs. We complemented these imaging assays with genetic and chemical perturbation experiments while measuring Pol II behavior in infected and uninfected cells. Finally, we performed ATAC-seq to sample the chromatin state of the viral DNA, and used Oligopaints to estimate the number of viral genomes within the RCs of infected cells.

Despite initial experiments showing that RCs display many of the macroscopic hallmarks of LLPS, we unexpectedly found that recruitment of Pol II and other DNA-binding proteins to RCs is achieved through a distinct compartmentalization mechanism not driven by IDR-dependent protein:protein interactions. Rather, we find that Pol II recruitment is achieved predominantly through transient, nonspecific binding of Pol II to viral DNA. These interactions occur independent of transcription initiation, and rely on the unusual feature of the HSV1 genome that it is largely maintained as “naked”, nucleosome-free DNA, which is much more accessible to DNA-binding proteins than host chromatin. Our findings show that nonspecific binding can play a key role in RC formation, in Pol II recruitment during infection, and more generally in the repertoire of distinct mechanisms a cell might employ to generate nuclear compartments.

## Results

### Pol II recruitment to Replication Compartments exhibits hallmarks of liquid-liquid demixing

HSV1 RCs form *de novo* following lytic infection, making them an attractive system to dissect compartment formation at the molecular level. To determine the mechanisms leading to the hijacking of Pol II, we used a U2OS cell line in which the catalytic subunit of Pol II has been fused to HaloTag (Boehning et al., 2018; Los et al., 2008). HSV1 infection occurs rapidly, with large replication compartments (RCs) forming within a few hours (Figure 1A). Because we were most interested in the early stages of lytic infection when Pol II is actively recruited to the RC, we focused our experiments on the period between 3 hours post infection (hpi) when RCs begin to emerge, and 6 hpi when infected cells begin to display significant cytopathic effects (Movies S1 and S2). To capture the earliest stages of Pol II recruitment, we used a low multiplicity of infection (MOI) to obtain a minimal dose of virions per cell.

**Figure 1.**
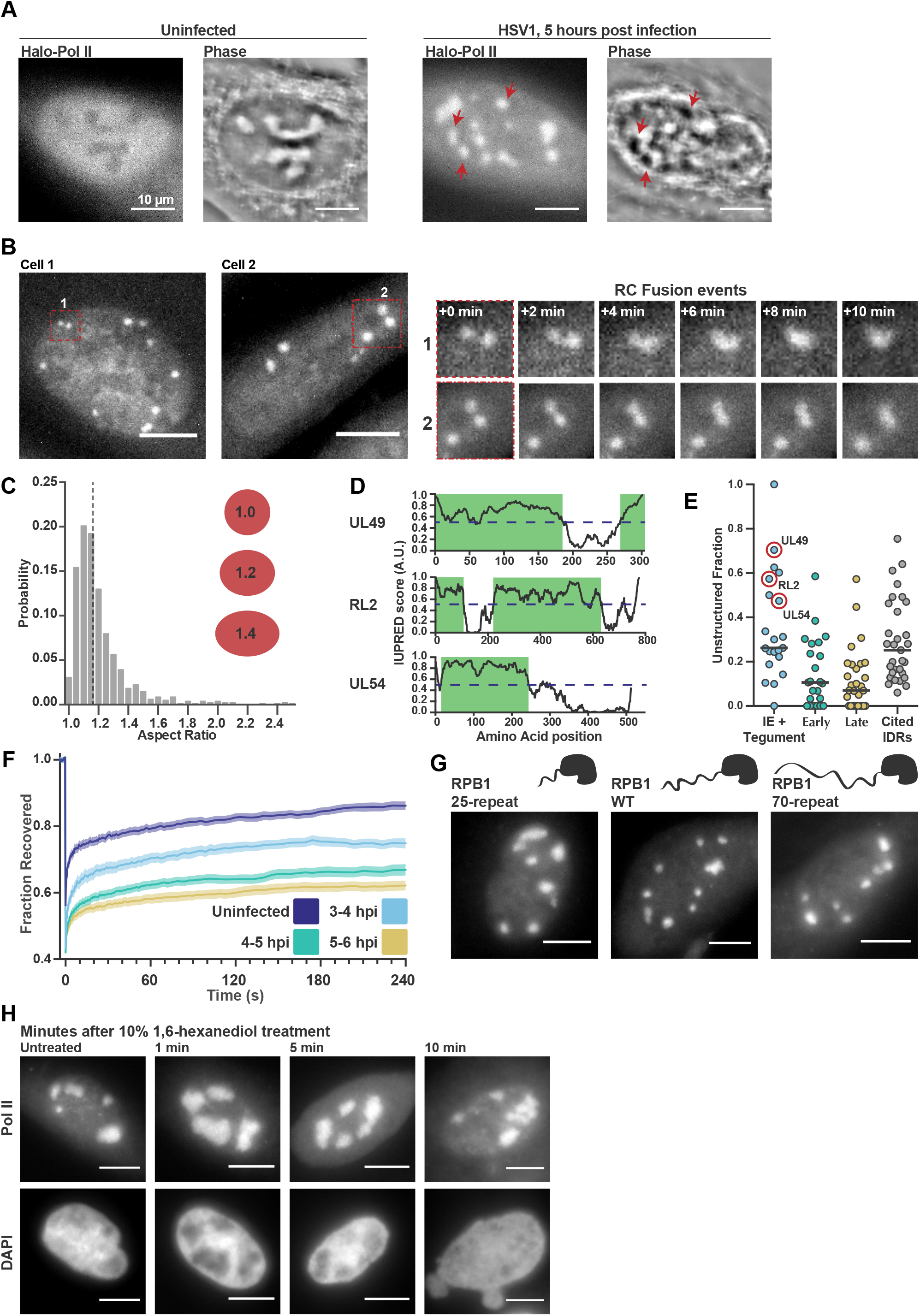
Pol II recruitment to Replication Compartments exhibit hallmarks of liquid-liquid demixing. **A)** Representative matched images of HaloTag-Pol II labeled with JF505 and Spatial Light Interference Microscopy to measure phase shifts for an uninfected cell, and for a cell 5 hpi. Arrows highlight examples of corresponding regions in the two images where an RC shows a significantly different phase value compared with the surrounding nucleoplasm. **B)** Time-lapse images from two cells (“Cell 1” and “Cell 2”). Zoom in shows RCs fusing beginning at t = 204 min and t = 220 min, respectively. HaloTag-Pol II U2OS cells were labeled with JF_549_ and infected, then imaged every 120 seconds beginning 3 hpi. See also Movies S1 and S2. **C)** A histogram of the aspect ratios (max diameter / min diameter) of RCs for 817 individual RCs from 134 cells, 3 to 6 hpi. The dotted line marks the median value of 1.18. Red ellipses provided a guide to the eye for different aspect ratios. **D)** IUPred scores for three viral proteins (UL49, RL2, UL54) as a function of residue position. Dashed line indicates an IUPred score of 0.5. Green boxes are predicted to be IDRs. **E)** The fraction of each protein in the viral proteome that is unstructured, separated by kinetic class. Immediate early (IE) and tegument proteins are as enriched in IDRs as a curated list of proteins containing IDRs known to drive phase separation (Cited IDRs). The proteins plotted in panel D are marked with a red circle. **F)** Fluorescence Recovery After Photobleaching (FRAP) curves of Pol II in RCs from 3-4 hpi, 4-5 hpi, and 5-6 hpi (n = 24, 33, and 33), compared with uninfected cells (n = 31). Curves represent the mean flanked by SEM. **G)** Representative images from cell lines expressing JF_549_-labeled HaloTag-RPB1 labeled with JF_549_ with a C-terminal domain containing different numbers of heptad repeats. **H)** Matched fluorescence images of JF_549_-labeled Pol II and DAPI. Cells were infected with HSV1 for 5 hours, then added 1,6-hexanediol to a final concentration of 10%, and fixed 1, 5 and 10 minutes after treatment. All scale bars are 10 μm.

In addition to Pol II, many other viral and nuclear factors re-localize to RCs (Dembowski and DeLuca, 2015). In fact, this redistribution of proteins is so dramatic that it can be seen by phase contrast microscopy as a change in the refractive index of RCs (Figure 1A). We observed, in agreement with previous studies, that RCs grow and move over the course of infection (Figure 1B) (Chang et al., 2011; Taylor et al., 2003). We also found that RCs exhibit several other behaviors characteristic of liquid droplets, such as an ability to fuse (Figure 1B, Movies S1 and S2) and a spherical shape, as indicated by an aspect ratio close to one (Figure 1C, median 1.18, n = 817). RCs are particularly round early in infection, when they are small (Figure S1). Such behaviors closely mimic the behavior of phase-separated liquid droplets, where the interface is thought to be subject to surface tension (Brangwynne et al., 2011; Feric et al., 2016).

Because LLPS is thought to be mostly driven by weak protein:protein interactions between intrinsically disordered regions (IDRs), we used the protein disordered region prediction algorithm IUPred to predict IDRs within viral proteins (IUPred > 0.55) (Dosztanyi et al., 2005; Dosztányi et al., 2005), and found these predictions match well with known disordered regions (Figure 1D) (Everett, 2000; Hew et al., 2015; Pfoh et al., 2015; Tunnicliffe et al., 2015). Across all viral proteins, we identified predicted IDRs longer than 10 amino acids, and used these to estimate what fraction of each protein sequence is unstructured (Figure 1E, Table S1). When categorized by temporal class, we noted that the immediate early (IE) and viral tegument proteins—the two groups that are presented to the cell first upon infection—had the highest fraction of predicted intrinsic disorder. In fact, when compared to a list of proteins known to undergo LLPS *in vitro*, the IE and tegument proteins are slightly more disordered (Figure 1E; Table S2 and citations within). Under the working hypothesis that multivalent interactions between protein IDRs drive phase separation, the similarity in predicted disorder profiles between this curated list and the IE and tegument proteins suggests that IDRs in viral proteins are as likely to drive LLPS as experimentally validated proteins.

Based on the above descriptive observations, we hypothesized that Pol II is recruited to RCs through interactions between its CTD and other IDR-containing proteins within the RC. To test this, we measured the FRAP dynamics of Pol II in RCs. We saw a consistent slowing of recovery as infection progressed and RCs got larger (Figure 1F), which could be interpreted as evidence that RCs act as a separate liquid phase that incorporates Pol II and sequesters it from the rest of the nucleoplasm. Subsequent experiments to directly test this hypothesis, however, cast doubt on this interpretation.

We recently reported that hub formation by Pol II in uninfected cells occurs in a manner dependent on the length of the Pol II CTD (Boehning et al., 2018). To test whether the Pol II CTD likewise mediates interaction with RCs, we compared Pol II accumulation in RCs in cells with the wild-type Pol II CTD (with 52 heptad repeats) and cell lines bearing truncated (25 repeats) or extended (70 repeats) CTDs. To our surprise, the length of the CTD had no detectable effect on its incorporation into RCs (Figure 1G), suggesting that Pol II does not require IDR interactions through its CTD to become enriched.

As a further test of the role of IDR interactions in Pol II accumulation in RCs, we treated cells with 1,6-hexanediol, which disrupts weak hydrophobic interactions between IDRs that drive LLPS (Boehning et al., 2018; Chong et al., 2018; Lin et al., 2016; Lu et al., 2018; Strom et al., 2017). We infected cells for five hours, and then subjected them to treatment with a high concentration (10% v/v) of 1,6-hexanediol for one, five, or ten minutes. Five minutes after treatment, the morphology of the nucleus began to change, and by ten minutes it was noticeably deformed, consistent with widespread disruption of cellular organization by 1,6-hexanediol (Lin et al., 2016). Nonetheless, Pol II remained highly enriched in RCs after 1,6-hexanediol treatment (Figure 1H), implying that formation of RCs does not require interactions between IDRs of Pol II and viral proteins.

### Pol II diffusion within and across RC boundaries is inconsistent with an LLPS model

The data outlined in Figure 1 present a potential contradiction, as RCs exhibit several properties commonly associated with phase separation *in vitro*, yet Pol II recruitment to RCs was not susceptible to disruption by 1,6-hexanediol or dependent on CTD length. Given these results, we sought to better understand the mechanism driving the enrichment of Pol II in RCs by measuring the behavior of individual Pol II molecules. To accurately capture both immobile and freely diffusing Pol II molecules, we used stroboscopic photo-activatable single particle tracking (spaSPT) to visualize and track molecules (Figure 2A) (Hansen et al., 2017, 2018). We labeled HaloTag-Pol II with equal amounts of JF_549_ and PA-JF_646_ (Grimm et al., 2015, 2016), which allowed us to monitor its overall distribution using the former dye while tracking individual molecules using the latter dye. We generated masks of the location of the nuclear periphery and individual RCs and used these masks to sort trajectories as either “inside” or “outside” of RCs (Figure 2B).

**Figure 2.**
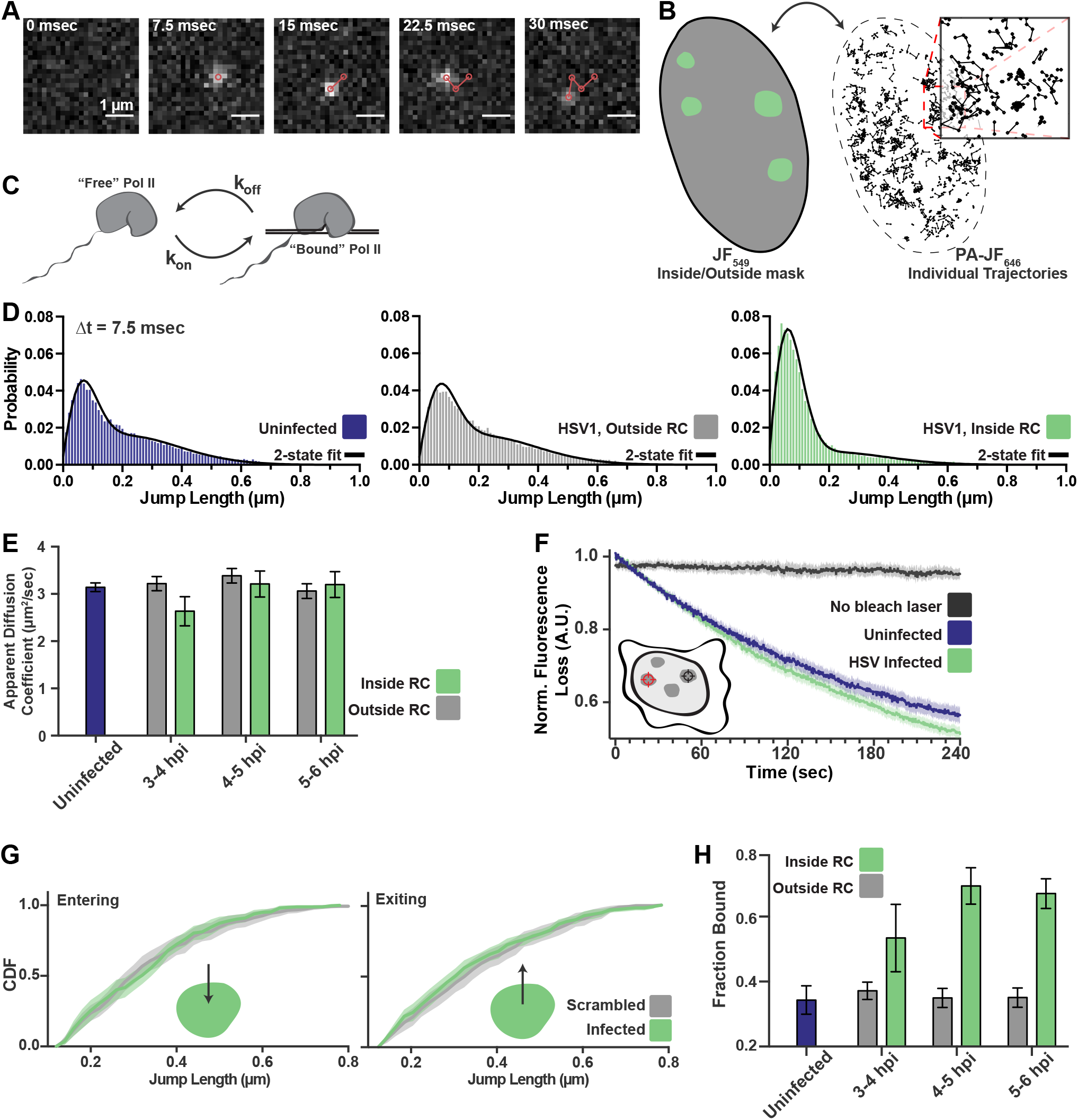
spaSPT of Pol II in infected cells shows no change in diffusion but an increase in binding. **A)** Example frames from a spaSPT movie, overlaid with the results from localization and tracking. Scale bar is 1 μm. **B)** Masks to identify the nuclear boundary (black line) and RCs (green regions) are generated with one color channel (JF_549_), and these annotations are applied to trajectories from the other color channel (PA-JF_646_). **C)** Depiction of two-state model where Pol II can either be freely diffusing, or DNA-bound, each with a characteristic rate constant. **D)** Jump length distributions between consecutive frames of spaSPT trajectories. Histograms pooled from uninfected cells (n = 27), or HSV1 infected cells between 4 and 6 hpi (n = 96). Each distribution is fit with a 2-state model. **E)** Mean apparent diffusion coefficient derived from the 2-state fit in (D). Error bars are the standard deviation of the mean, calculated from 100 iterations of randomly subsampling 15 cells without replacement and fitting with the model. **F)** Fluorescence Loss In Photobleaching (FLIP) curves comparing the rate of fluorescence loss after photobleaching JF_549_-labeled Pol II in uninfected cells and HSV1 infected cells. A 1 μm bleach spot is placed inside an RC (red crosshairs) and bleached between every frame. The loss of fluorescence is measured from another RC (black crosshairs). **G)** Cumulative Distribution Function (CDF) of the mean flanked by the SEM for jump lengths of molecules entering (left) or exiting (right) RCs. Distribution for HSV1 infected cells is compared to the distribution of molecules entering/exiting compartments that have been randomly shuffled around the nucleus *in silico*. **H)** Mean fraction of bound molecules derived from the 2-state fit in (D). Error bars are the standard deviation of the mean, calculated from 100 iterations of randomly subsampling 15 cells without replacement and fitting with the model. See also Figures S2, S3A-D, and S4.

Quantitative measurements can be made by building histograms of all the displacement distances from the trajectories and fitting to a two-state model in which Pol II can either be freely diffusing (“free”), or immobile and hence presumably bound to DNA (“bound”) (Figure 2C). Such a two-state model gives two important pieces of information: the fraction of “bound” and “free” molecules, and the apparent diffusion coefficient of each population (Hansen et al., 2018). It is important to note that, because this modeling approach takes the aggregate of many thousands of molecules, these data cannot measure how long a particular molecule remains bound in a given binding event. Here, “bound” refers to both specific DNA binding events—e.g. molecules assembled at a promoter or engaged in mRNA elongation—as well as transient, nonspecific binding interactions.

The difference in the behavior of Pol II inside of RCs compared with the rest of the nucleoplasm is immediately apparent from examining the lengths of jumps between consecutive frames (Figure 2D). Fitting a two-state model to the data, we were surprised to find that the mean apparent diffusion coefficient of the free population was unchanged between trajectories inside of RCs compared with those outside RCs or in uninfected cells. If RCs were a *bona fide* separate phase, one would expect differences in molecular crowding or intermolecular interactions to predominantly affect free diffusion, resulting in substantially different diffusion coefficients between the populations (Bergeron-Sandoval et al., 2016). Furthermore, the similarity in diffusion coefficients between infected and uninfected cells argues against a separate viral protein-Pol II complex responsible for recruitment to RCs.

We verified this result in two ways: First, we performed a fluorescence loss in photobleaching (FLIP) experiment, in which a strong bleaching laser targets the inside of an RC and loss of fluorescence elsewhere in the nucleus is measured to quantify exchange of Pol II between the nucleoplasm and the RC. Consistent with the spaSPT data, we see that Pol II molecules exchange between RCs and the rest of the nucleoplasm as fast, if not faster, than Pol II in an uninfected cell (Figure 2F). Similar results were obtained by using Pol II tagged with the photo-convertible fluorescent protein Dendra2 (Cisse et al., 2013) and photo-converting, rather than bleaching, molecules in the RC (Figure S2). Thus, Pol II molecules diffuse out of the RC, rather than remaining sequestered within a single compartment. Second, a liquid-liquid phase separation model predicts that a diffusing Pol II molecule within an RC will be more likely to remain within the RC than to exit the RC when it reaches the compartment boundary. To test this prediction, we examined all trajectories for events in which a molecule crosses from inside of the RC to outside, or vice versa, to look for evidence of such a constraint. Comparing the distribution of displacements for a particle going from inside the RC to outside, we see no difference in the distribution of displacements, either entering or leaving RCs, when compared to uninfected cells in which mock RC annotations were randomly imposed *in silico* (Figure 2G, Figure S3). With these experiments we cannot detect any evidence of a boundary for molecules entering or leaving RCs, further arguing that this compartment does not consist of a distinct liquid phase.

While the two-state model shows no change in diffusion coefficient of molecules inside versus outside RCs, the fraction of molecules in the “bound” state nearly doubles inside RCs, reaching ~70% (Figure 2H). The diffusion coefficients we measured with the bound populations are still consistent with those of chromatin (Hansen et al., 2018), indicative that these populations reflect DNA binding (Figure S4). The increase in the fraction of bound molecules is further supported by the FRAP data (Figure 1F). We verified this was not an artifact of the masking process by using the same process of artificially imposing RCs randomly *in silico* (Figure S3), and we found no difference in the fraction of bound molecules. Such a significant shift in the relative populations of bound and free molecules inside RCs, taken together with the previous data, shows that the mechanism driving Pol II recruitment to RCs is dominated by DNA binding, rather than by IDR-mediated interactions that sequester Pol II away in a separate liquid phase.

### Pol II recruitment to RCs occurs independent of transcription initiation

The above data argue against formation of RCs by LLPS, suggesting that some other mechanism must explain the doubling of DNA-bound Pol II in RCs. One possibility is that increased recruitment of Pol II is explained by high levels of active transcription within RCs. Multiple lines of evidence suggest that transcription derived from the viral genome is activated to a much greater extent than even the most highly transcribed host mRNA (Rutkowski et al., 2015) and so an enriched population of actively elongating Pol II would be expected to increase the “bound” population.

To test whether active transcription is necessary for Pol II recruitment to RCs, we treated infected cells with either Flavopiridol or Triptolide, two small molecules that selectively inhibit different stages of transcription initiation (Figure 3A). Flavopiridol is a potent inhibitor of CDK9 that prevents the phosphorylation of serine-2 of the RPB1 CTD heptad, and thus prevents transcription from proceeding beyond ~50 nucleotides downstream of the transcription start site (TSS) (Chao and Price, 2001; Jonkers et al., 2014). Triptolide, on the other hand, is an inhibitor of the ATPase domain of TFIIH, preventing the stable engagement of Pol II at the Pre-Initiation Complex (PIC) (Alekseev et al., 2017; Chen et al., 2015; Titov et al., 2011). Samples treated with Triptolide show a loss of engaged Pol II at promoters (Jonkers et al., 2014; Shao and Zeitlinger, 2017).

**Figure 3.**
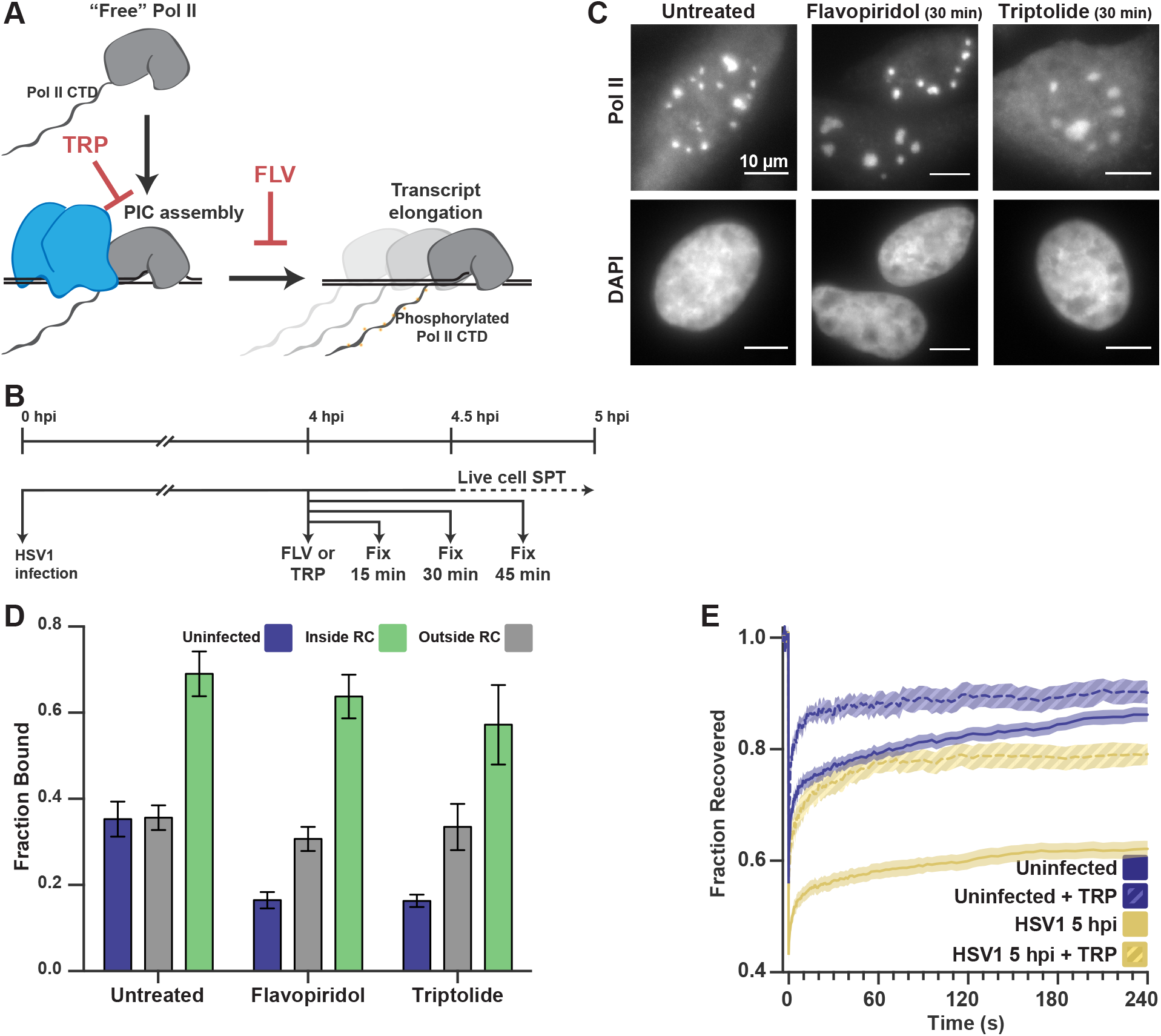
Pol II recruitment to RCs occurs independent of active transcription. **A)** Schematic of Pol II-mediated transcription inhibition. Triptolide (TRP) prevents stable Pol II engagement with the assembled pre-initiation complex. Flavopiridol (FLV) inhibits engaged Pol II from elongating past ~50 bp. B) Schematic of experiment regimen for imaging infected cells after transcription inhibition. Cells are infected at time 0. At 4 hpi, cells are treated with transcription inhibitors Flavopiridol (FLV) or Triptolide (TRP) and fixed after 15, 30, or 45 minutes. For live cell imaging, cells are treated with drug at 4 hpi, and after 30 minutes, they are mounted on the microscope and imaged directly. C) Representative images of JF_549_-labeled HaloTag-Pol II and DAPI after 45 minutes of Triptolide or Flavopiridol treatment. All scale bars are 10 μm. D) Mean fraction bound measured from spaSPT of HaloTag-Pol II, after either Flavopiridol or Triptolide treatment. Error bars are the standard deviation of the mean, calculated from 100 iterations of randomly subsampling 15 cells without replacement and fitting with the model. E) FRAP recovery curves of Pol II with (hashed) and without (solid) Triptolide treatment, for uninfected cells (N = 31, 9 respectively) and cells infected with HSV1, 5hpi (N = 32, 12 respectively). Also see Figure S5.

HSV1 RCs require the expression of immediate-early and early genes to generate the DNA replication machinery, so we allowed the infection to progress for four hours before treating with either compound. Cells at this timepoint have well formed RCs, and Pol II binding is already greatly increased (Figure 2H). We treated these cells with 1 μM Flavopiridol or 1 μM Triptolide for 15, 30, or 45 minutes to allow any elongating polymerases to finish transcribing (Figure 3B). After treatment, we fixed cells, and we performed RNA fluorescence in situ hybridization (FISH) using a probe against an intronic region to detect nascent transcripts and, in parallel, and immunofluorescence to mark the DNA-binding protein ICP8, a common marker for RCs (Taylor et al., 2003). After 30 minutes of drug treatment, transcription is significantly reduced (Figure S5). Remarkably, even after 45 minutes of treatment, ~80% of the Pol II signal remains within RCs (Figure 3C, Figure S5). These data suggest that the recruitment of Pol II to RCs occurs largely independently of transcription, or even stable engagement with gene promoters.

We next tested whether treatment with these transcription inhibitors would change the bound fraction measured by spaSPT. In uninfected cells, Triptolide or Flavopiridol treatment both reduce the fraction of bound Pol II by half, to ~15% (Figure 3D), similar to what others have reported (Boehning et al., 2018; Teves et al., 2018). Surprisingly, inhibition of transcription with Flavopiridol reduced the bound fraction inside of RCs by only ~5% (Figure 3D). Even treatment with Triptolide, which prevents stable engagement with TSS-proximal DNA only reduced the fraction bound by ~12% (Figure 3D). We were surprised to see that with either drug treatment, HSV1 infection appears to also confer some resistance to the effects of the drugs on Pol II binding to host chromatin, despite the fact that these concentrations of transcription inhibitors are sufficient to prevent new transcription (Figure 3D, Figure S5). Given the inherent limitation of spaSPT for inferring the length of binding events, we wanted to confirm that drug treatment prevented stable Pol II binding. Indeed, FRAP experiments in infected cells treated with Triptolide show a dramatically faster recovery rate (Figure 2E). For the infected samples, this means that the “bound” molecules measured by SPT do not remain bound for long times, as one would expect from high affinity protein-protein or protein-DNA interactions at cognate sites. Instead, these binding events are likely short compared to the timescales of PIC assembly and transcription initiation. Such highly transient binding events also argue against sequence specific, high affinity interactions as drivers for sequestering Pol II to the RC. The fact that infected cells show resistance, in terms of DNA binding, to the drug treatment may be a result of other viral mechanisms that occur during infection, such as aberrant Pol II CTD phosphorylation (Rice et al., 1994) or termination defects (Rutkowski et al., 2015). Still, our results suggest that viral DNA and/or DNA-associated proteins mediate very rapid, predominantly nonspecific, interactions within RCs.

### HSV1 DNA is more accessible than host chromatin to Pol II

The result that Pol II molecules remain bound—however transiently—to the viral DNA, even in the absence of transcription, suggests that the DNA itself likely plays a dominant role in Pol II enrichment in RCs. To our knowledge, the genome copy number present in individual RCs has never been determined, but this information is crucial to understand the role viral DNA may play in RC formation and function. We therefore sought to measure the amount of DNA in RCs using Oligopaints, a variant of DNA fluorescence in situ hybridization, to target fluorescent probes to two specific regions of the viral genome (Figure 4A) (Beliveau et al., 2012; Boettiger et al., 2016). Infected cells were fixed three, four, five, and six hours post infection. The amount of DNA was measured by fluorescence intensity of the compartment, determined independently for the two probe sets. These fluorescence intensities were compared to samples that were infected in the presence of phosphonoacetic acid (PAA), a compound that prevents replication of the viral DNA and thus ensures that there is one copy of the viral genome per punctum (Figure 4B) (Eriksson and Schinazi, 1989).

**Figure 4.**
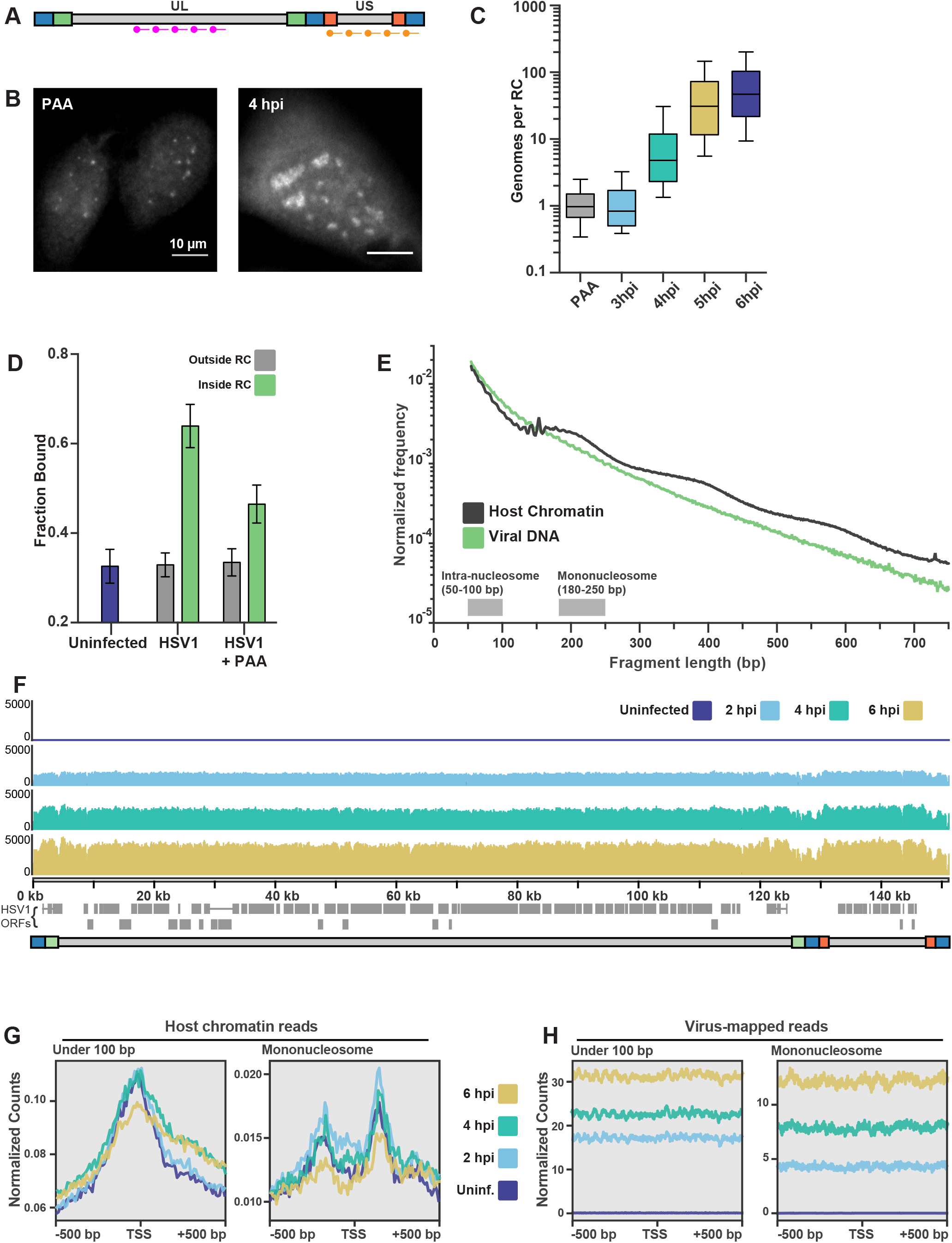
ATAC-seq reveals HSV1 DNA is much more accessible than chromatin. **A)** Schematic representation of where Oligopaint probes target within the viral genome for DNA Fluorescence In Situ Hybridization. AlexaFluor 647-labeled probes target a 10,016 bp region in the middle of the Unique Long (UL) arm. AlexaFluor 555-labeled probes target a 7703 bp region in the middle of the Unique Short (US) arm. **B)** Representative images of DNA FISH on cells infected in the presence of the replication inhibitor phosphonoacetic acid (PAA, left) and 4 hpi (right). Pixel intensity values are the same for the two images. Scale bars are 10 μm. **C)** Quantification of fluorescence intensity of DNA FISH signal in RCs at increasing times after infection. Data are normalized to the median intensity value of cells infected in the presence of the replication inhibitor PAA. Medians are indicated by a solid line (PAA = 1, 3 hpi = 0.83, 4 hpi = 4.77, 5 hpi = 31.13, 6 hpi = 46.95). Also see Figure S5 **D)** Mean fraction bound extracted from 2-state model of PA-JF_646_-labeled HaloTag-Pol II, comparing HSV1 infected cells between 4 and 6 hpi to cells infected in the presence of PAA. Error bars are the standard deviation of the mean, calculated from 100 iterations of randomly subsampling 15 cells without replacement and fitting with the model. **E)** Fragment length distribution of ATAC-seq data for cells 4 hpi. Reads mapping to the human genome are in gray, and reads mapping to the HSV1 genome are in green, and data are normalized to the total number of reads mapping to each organism. Lengths corresponding to intra-nucleosomal DNA (50–100 bp) and mononucleosomal DNA (180-250 bp) are marked as a reference. **F)** ATAC-seq read density for all fragment lengths plotted across HSV1 genomic coordinates for uninfected cells, and for 2, 4, and 6 hpi. **G)** ATAC-seq analysis of intra-nucleosomal DNA (50–100 bp) and mononucleosomal DNA (180-250 bp). Global analysis of all human Pol II-transcribed genes, centered at the transcription start site (TSS) at different times after infection. **H)** The same analysis as in (G), but centered at the TSS of HSV1 genes. Also see Figures S6 and S7.

While there is a great deal of expected RC-to-RC heterogeneity, the number of genomes within an RC correlates well with the time post infection (Figure 4C). There is also a strong correlation between RC size and genome copy number (Figure S6). Based on these data, we calculate that the average RC at 6 hpi has a DNA concentration of 3.9 ×10^4^ bp/μm^3^, approximately 240 times less concentrated than average host chromatin. The sum of all RCs in an average infected cell corresponds to just ~0.2% of total DNA in karyotypically normal human nuclei (Table S3). Despite being orders of magnitude lower in DNA content and concentration, inhibition of viral DNA replication with PAA caused a ~20% decrease in the fraction of bound Pol II molecules inside the pre-replication foci as measured by spaSPT, nearly down to the level of host chromatin (Figure 4D). This, despite the fact that all immediate early and early genes, including many of the proteins known to interact with Pol II, are still highly expressed in PAA-treated samples (Lester and DeLuca, 2011; Zhou and Knipe, 2002).

Since most of the Pol II binding events that we observe inside of an RC appear to be unrelated to transcription, but clearly dependent on viral DNA replication, we wondered what might be different about the viral genome relative to host chromosomes. A likely candidate is the chromatin state of the viral DNA. There is presently no clear consensus about the organization of viral DNA during lytic infection. Multiple studies have successfully used ChIP to sample histone marks to determine histone association with the viral DNA (Bloom et al., 2010; Lang et al., 2017; Lee et al., 2016), though this may also be explained by formation of nonnucleosomal histone-DNA intermediates (Torigoe et al., 2011), especially since mass spectrometry studies have failed to detect histones associated with viral DNA (Dembowski and DeLuca, 2015; Taylor and Knipe, 2004). Moreover, immunofluorescence against histones shows no detectable signal in RCs (Dembowski and DeLuca, 2015). In addition, one function of viral ICP0 is to actively evict histones from DNA, which suggests that the HSV1 genome is maintained largely free of histones (Lee et al., 2016).

To test histone occupancy of the viral DNA, and get a measure of its accessibility, we turned to ATAC-seq, which gives signal proportional to the accessibility of the DNA at a given locus (Buenrostro et al., 2013). We infected our HaloTag-Pol II cell line, and performed Tn5 transposition reactions at 2, 4, and 6 hpi. We also included a sample that was uninfected, and one infected in the presence of PAA. At all times after infection, the distribution of fragment lengths mapping to the viral genome showed a much faster decay, and no evidence of nucleosomal laddering, in contrast to reads that map to the host genome (Figure 4E, Figure S7). When we visualized the reads along the viral genome, the profiles were strikingly flat and featureless, suggesting that all regions of the viral genome are equally accessible to Tn5 (Figure 4F).

Based on the amount of viral DNA present in an infected cell, we calculated the fraction of reads one would expect to map to the virus relative to the host. At 6 hpi, under our infection conditions, viral DNA represents an average 0.2% of total nuclear DNA content. Yet, at this time point, 24.2% of reads mapped to the virus on average. From this, we calculate that DNA inside of RCs is two orders of magnitude more accessible, despite its overall lower DNA concentration relative to host DNA (Table S3).

In metazoan genomes, active genes can be identified by their high accessibility (Thurman et al., 2012). An average of all annotated human mRNA genes, centered at the TSS, shows a characteristic peak of accessibility at the TSS for reads with a length corresponding to inter-nucleosomal distances (<100 bp), and a characteristic trough of mononucleosome sized fragments (180 – 250 bp) (Figure 4G). By contrast, TSS averages mapped to the viral genome for either short or mono-nucleosome fragments show no changes in accessibility. Thus, even averaging over all viral transcripts, it is clear that the entire viral DNA remains equally accessible (Figure 4H). Taken together, these data indicate that the HSV genome is maintained in a largely nucleosome-free state and thus, highly accessible to DNA binding proteins such as Pol II.

### Transient DNA-protein interactions drive Pol II hub formation through repetitive exploration of the replication compartment

Knowing that the DNA inside RCs is vastly more accessible to nuclear factors than the host chromatin, we next asked what emergent properties of this accessible DNA might help explain Pol II recruitment. We took advantage of a viral strain that is able to incorporate chemically modified nucleotides during replication (Dembowski and DeLuca, 2015), to label newly replicated viral DNA with Alexa Fluor 647, and thus allow DNA in the RCs to be visualized at high resolution using stochastic optical reconstruction microscopy (STORM) (Figure 5A) (Rust et al., 2006). Unlike host chromatin, whose overall density and compaction scales reproducibly with domain size for active chromatin (Boettiger et al., 2016), viral DNA shows a spatial variability in local density of nearly three orders of magnitude.

**Figure 5.**
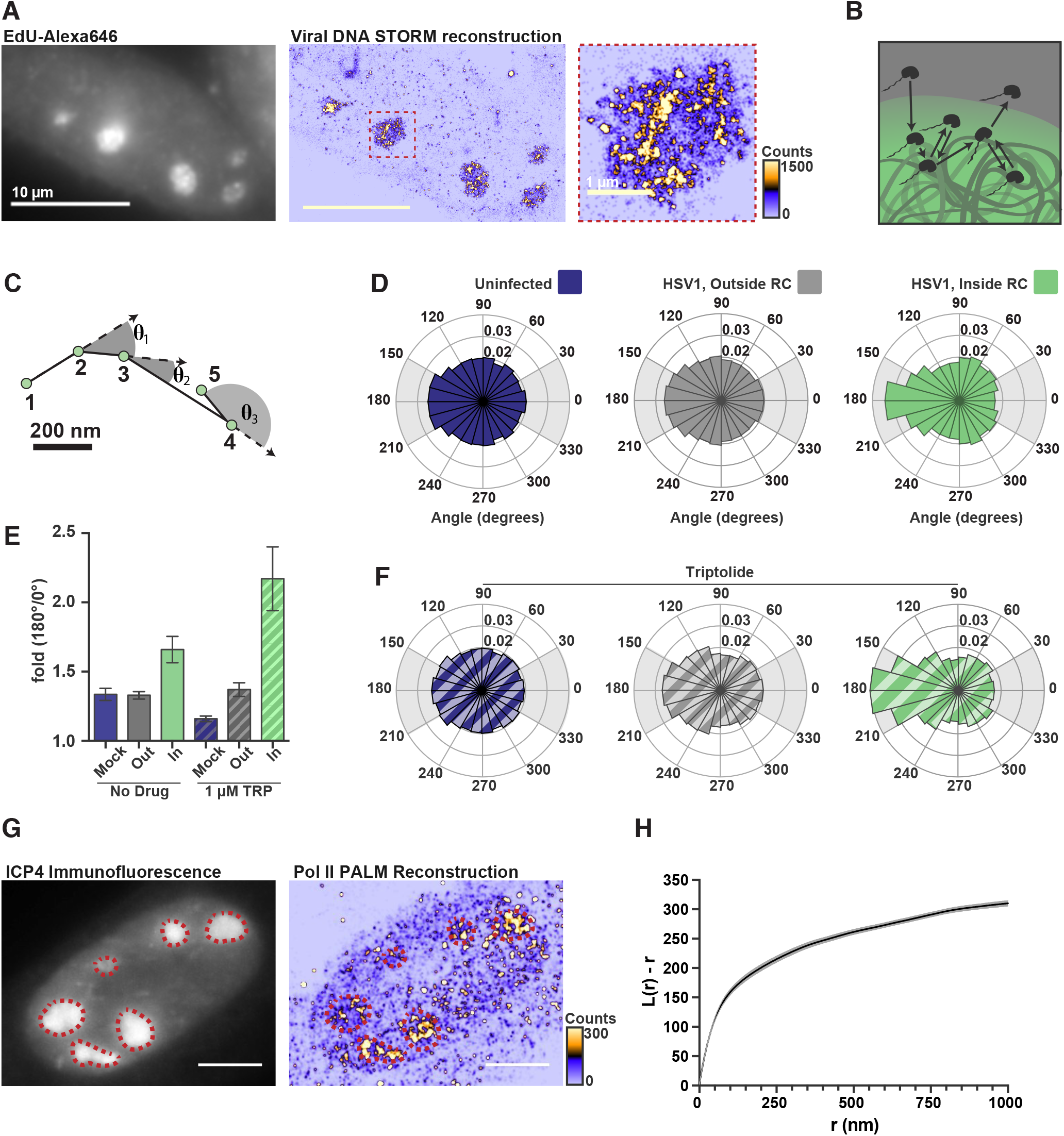
DNA-binding alters Pol II exploration of RCs. **A)** Representative STORM image of an HSV1 infected cell in the presence of EdU to allow labeling with AlexaFluor 647 and matched diffraction-limited images and STORM reconstruction, rendered with 20 nm by 20 nm pixels. Zoom-in shows one RC, and the heatmap shows the number of fluorophore localizations occurred in each rendered pixel. **B)** Schematic of Pol II entering an RC and randomly sampling the viral DNA, sometimes revisiting the same, or closely adjacent, sites before diffusing away. **C)** Example spaSPT trace, with each set of three localizations (n, n+1, n+2) forming a measurable angle (θ_n_…). **D)** Angular distribution histograms extracted from diffusing HaloTag-Pol II molecules in uninfected cells, and HSV1 infected cells 4-6 hpi, inside and outside of RCs. Shaded regions highlight the angles used to calculate the relative probability of moving backward compared to forward in (E). **E)** Quantification of the relative probability of moving backward compared to forward (180° ± 30° / 0° ± 30°). Error bars are the standard deviation of the mean, calculated from 100 iterations of randomly subsampling 15 cells without replacement. **F)** Same as in (D), except that cells were treated with Triptolide at least 30 minutes prior to imaging. Quantification of this data is also show in (E). **G)** Representative PALM image of PA-JF_646_-labeled HaloTag-Pol II, rendered with a pixel side of 20 nm by 20 nm. Infected cells were identified by immunofluorescence against the HSV1 protein ICP4. Heatmap corresponds to the number of detections per rendered pixel. Dotted red lines mark the outlines of RCs, as identified by the immunofluorescence image. **H)** L-modified Ripley Curve (L(r)-r) for HaloTag-Pol II inside of RCs in cells 5 hpi (n = 13 cells). Graph shows the mean flanked by the SEM. All scale bars are 10 μm. Also see Figure S3E and S3F.

The greater accessibility and higher variability in local density of viral DNA lend themselves to a possible mechanism by which Pol II becomes enriched. Recent theoretical work has shown that a polymer, like DNA, which has many binding sites in close proximity can induce a particle to revisit the same or adjacent sites repetitively during its exploration of the nucleus (Amitai, 2018) (Figure 5B). In such a case, we should be able to see signatures in our spaSPT dataset of Pol II continually revisiting adjacent sites on the viral DNA. To check, we calculated the angle formed by every three consecutive displacements and compiled these angles into a histogram for all particles strictly identified as freely diffusing (see Methods) (Figure 5C) (Izeddin et al., 2014). For particles experiencing ideal Brownian motion, the angular histogram will be isotropic. Anisotropy can arise either by imposing reflective boundaries on the particle, or adding the aforementioned “traps” thereby giving the particle a greater probability of revisiting proximal sites before diffusing away (Amitai, 2018).

In uninfected cells, and in infected cells outside of RCs, Pol II displays diffusion that is only mildly anisotropic, consistent with mostly Brownian motion throughout the nucleus. In stark contrast, inside RCs Pol II diffusion is more anisotropic, especially around 180° (Figure 5D). To compare across samples, we computed the likelihood of a backward translocation (180° ± 30°) relative to the likelihood of a forward translocation (0° ± 30°). Analyzed this way, Pol II in an uninfected nucleus has a 1.3-fold greater chance of moving backward after a given translocation than it has of moving forward (Figure 5E). As expected, Pol II outside of RCs in infected cells has a nearly identical value to Pol II in an uninfected cell. Inside of an RC, however, that probability increases to 1.7-fold, showing that this effect is unique to RCs (Figure 5E). In cells treated with Triptolide, we see that when stable binding is inhibited, the effect created by transient binding events is further amplified (Figure 5E). Under this condition, Pol II inside an RC is 2-fold more likely to have a backward displacement after a forward one (Figure 5D), which helps explain the dramatic retention of Pol II inside RCs, even 45 minutes after inhibition of transcription (Figure 3C). Importantly, in uninfected cells where RC annotations have been shuffled *in silico*, no additional anisotropy is observed (Figure S3).

These data are most consistent with a model in which Pol II repetitively visits the highly accessible viral genome via multiple weak, transient binding events that result in Pol II rapidly hopping along the DNA. The sharp anisotropy of the molecular exploration within the RC means that a given Pol II molecule that enters an RC is more likely to visit the same site, or sites close in proximity, multiple times before it either finds a stable binding site or diffuses away.

The heterogeneous distribution of viral DNA within RCs, and the anisotropic way Pol II explores RCs, is also borne out in the distribution of Pol II molecules. We performed 3D photoactivated localization microscopy (3D PALM) on cells, using adaptive optics for precise 3D localization of individual Pol II molecules (Betzig et al., 2006; Izeddin et al., 2012). Similar to the viral DNA, PALM renderings of infected nuclei revealed a heterogeneous Pol II distribution within RCs (Figure 5G). For a more quantitative determination of Pol II clustering, we used Ripley’s L-function, a measure of how a spatial point pattern deviates from randomness (Figure 5H) (Nicovich et al., 2017). Here, a value greater than zero indicates a concentration of points higher than predicted for complete randomness at that given radius. For very small radii, a high L(r)-r value is likely due to blinking and other photo-physical artifacts (Annibale et al., 2011). However, our measurements of Ripley’s L-function remains well above zero and increases for all radii between 0 and 1000 nanometers, suggesting that Pol II forms hubs within RCs and that this clustering occurs at multiple length scales. This is consistent with other recent studies of Pol II in uninfected cells (Boehning et al., 2018), and in contrast to a structural protein like the CCCTC-binding factor (CTCF), whose L(r)-r curve shows clusters of a single characteristic size (Hansen et al., 2017).

### Nonspecific interactions with viral DNA license recruitment of other proteins

Seeing that Pol II is recruited to RCs via transient and nonspecific binding to the viral genome made us wonder whether this effect was specific to Pol II, or whether DNA accessibility can generally drive the recruitment of DNA-binding proteins to RCs. Certainly, many other DNA-binding proteins are recruited to RCs (Dembowski and DeLuca, 2015; Taylor and Knipe, 2004). To assess whether nonspecific DNA binding could be responsible for accumulation of other proteins within RCs, we looked at enrichment of the tetracycline repressor (TetR). TetR is a sequence-specific transcription factor found in bacteria that binds with high affinity to the 19 bp *tetO* sequence, which is absent in both human and HSV1 genomes (Bolintineanu et al., 2014). Thus, we reasoned that if nonspecific DNA association is the mechanism driving recruitment to the RC, TetR should also be recruited to the RC.

We transiently transfected TetR-GFP into the HaloTag-Pol II cell line, then infected them with HSV1 the following day. TetR-GFP, lacking a nuclear localization signal (NLS), ubiquitously occupies both the nucleus and cytoplasm. As predicted by our model, GFP signal was enriched inside of RCs (Figure 6). Pixel line scans of the matched Pol II and TetR channels show that the level of enrichment is modest (~25% over background for TetR, compared with ~200% for Pol II), but the two signals showed a high Spearman correlation (r > 0.77). A fluorescent protein with only an NLS showed no enrichment at RCs in infected cells (Figure S8). Thus, even a sequence-specific transcription factor with no cognate binding sites in the viral genome can be recruited to RCs based on its modest affinity for nonspecific DNA sequences.

**Figure 6.**
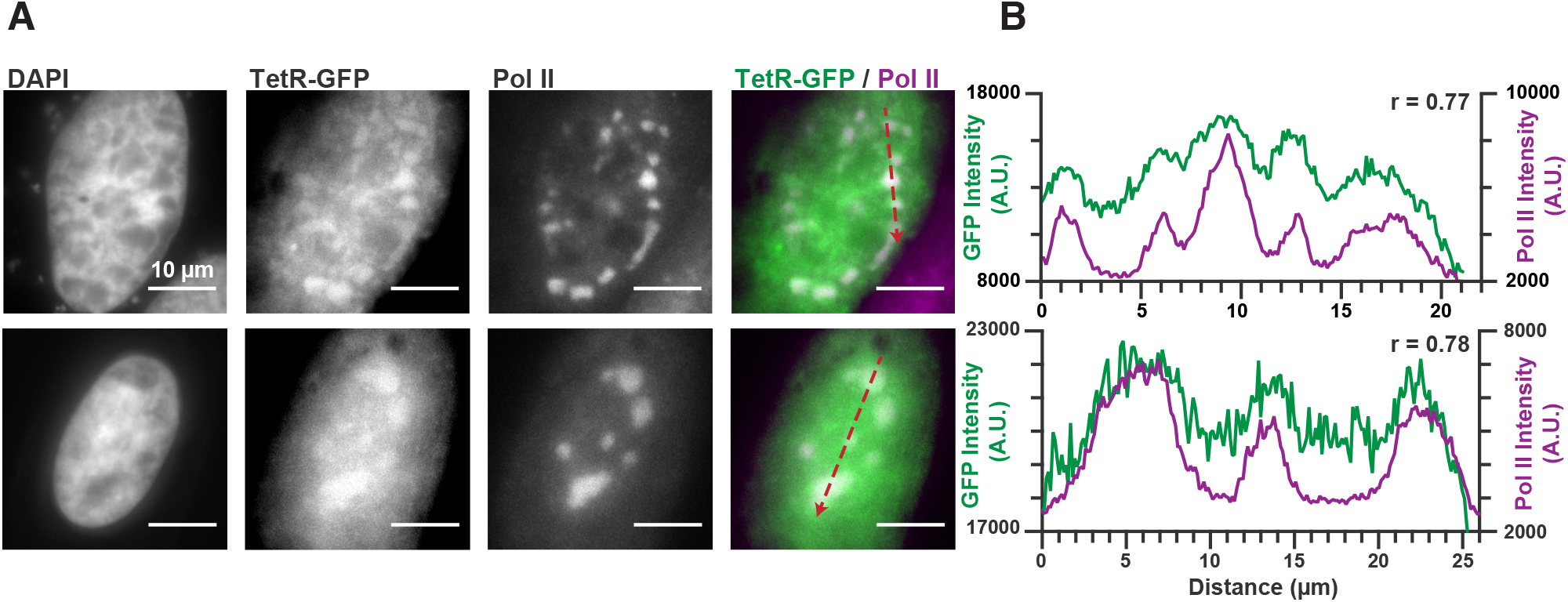
Nonspecific DNA binding drives accumulation of other factors in RCs. **A)** Representative images of HaloTag-Pol II cells transiently transfected with TetR-GFP prior to infection with HSV1. **B)** Pixel line scans of images in (A). Red arrows in (A) give the direction of the x-axis. Left axis is the intensity of TetR-GFP fluorescence, right axis is the intensity of JF_549_-labeled HaloTag-Pol II fluorescence. Spearman’s correlation was used to measure the covariation in the two channels across the line. All scale bars are 10 μm. Also see Figure S8.

These data suggest a model in which viral Pol II recruitment consists of transient, nonspecific binding/scanning events of the highly exposed viral genome (Figure 7A). A DNA-binding protein exploring the nucleus (uninfected, or infected but outside of RCs) may encounter some occasions for nonspecific interaction with duplex DNA, but because of the condensed nature of the host chromatin, these binding/scanning events are necessarily short (Figure 7B). In addition, it may take a protein many thousands of these transient binding events to finally reach a high-affinity site (Normanno et al., 2015). Within RCs, multiple copies of the highly accessible HSV1 genome are present, nonspecific events happen more frequently, with fewer and shorter 3D excursions between DNA contacts (Figure 7C), leading to redundant exploration of the RC and local accumulations of protein. This enrichment becomes even more skewed when aided by a high density of other interactors—for example when transcription is active on the viral DNA, and thus the both specific and general transcription factors are enriched via the same mechanism. Out data strongly indicate, though, that host Pol II accumulation at HSV1 RCs is not dependent on active transcription but instead is largely driven by transient, nonspecific protein-DNA interactions resulting from the high accessibility of the viral DNA.

**Figure 7.**
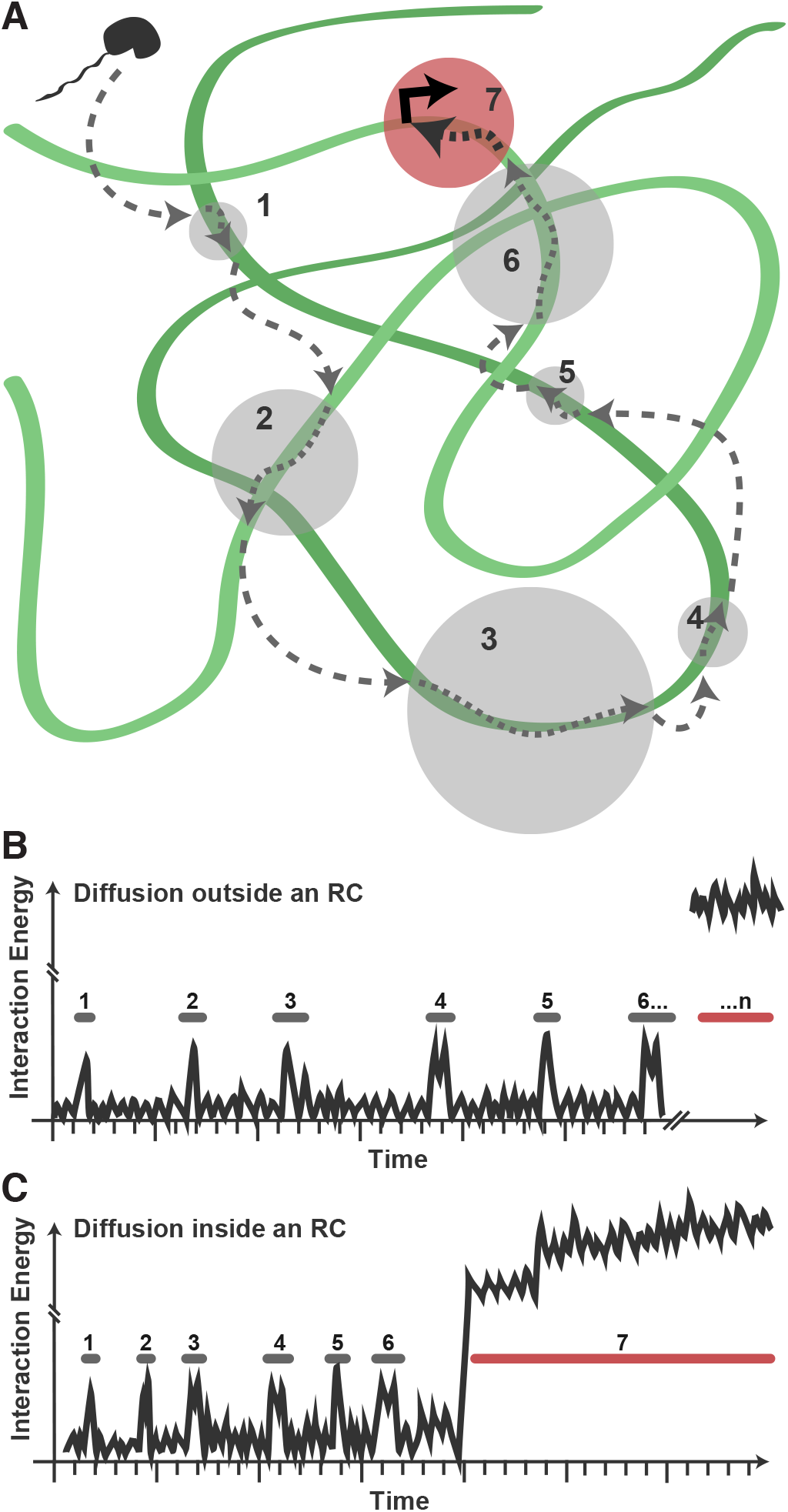
Model for Pol II exploration of RCs. **A)** A Pol II molecule encounters the accessible viral DNA multiple times along one potential route to eventually bind at a promoter. 3D diffusion through the RC is interrupted by binding interactions with the viral DNA (gray circles). These binding interactions may be very transient in nature, and may involve direct interaction with the viral DNA, 1D scanning along the DNA, or transient interactions with other DNA-associated proteins. **B)** Hypothetical comparison of nuclear exploration outside RCs as a function of time and binding energy. A DNA-binding protein in the chromatinized nucleus will encounter naked DNA sporadically, making multiple low-affinity interactions before eventually finding a high affinity site. **C)** Inside an RC, the high DNA accessibility might shorten the length of 3D excursions before a DNA-binding protein encounters another region of viral DNA in a low affinity, nonspecific interaction. This, in turn, may reduce the distance a molecule might diffuse before its next binding event, and increases both the chances of that molecule remaining in close proximity and the chances that it will find a high binding energy interaction.

## Discussion

### Multiple routes to create high local concentrations

Here we have demonstrated that Herpes Simplex Virus type 1 accumulates Pol II in replication compartments through a novel mechanism: its unusually accessible DNA genome provides many potential nonspecific binding sites, which causes a net accumulation of Pol II, acting as a molecular sink even in the absence of transcription. Such a mechanism for locally concentrating proteins is revealing, as it neither requires the formation of a stable macromolecular structure nor produces any behaviors at the single-molecule level suggesting a separate liquid phase. It is particularly striking because, from the macroscopic view, Pol II recruitment to RCs appears to share many of the behaviors commonly attributed to liquid-liquid phase separation—enrichment of proteins that have high intrinsic disorder, spherical, dynamic structures that undergo fusion, and a change in refractive index—and yet RCs are clearly a distinct class of membraneless compartments.

Given the high prevalence of IDRs in the viral proteome, it is likely that they have crucial functions in other aspects of the viral lifecycle, such as the assembly of the capsid or packaging of tegument proteins prior to envelopment. It still remains a possibility that these IDRs form some phase-like system inside of RCs. Crucially, our data demonstrate that, even if this is the case, it does not contribute to the enrichment or entrapment of Pol II. Our results prompt the need for a better characterization of *bona fide* phase separation, with a focus on its functional consequences *in vivo*. These data underscore the importance of rigorously dissecting the diverse mechanisms driving subcellular compartment formation, and suggest that caution should be exercised before immediately assigning LLPS as the primary assembly mechanism or interpreting the functional role of a phase-separated system solely based on macroscopic behaviors.

We emphasize that this is certainly not the only means by which herpesviruses interact with Pol II; many other studies have carefully documented the roles of both host and viral proteins in recruiting Pol II to transcribe viral genes (Davis et al., 2015; Zhou and Knipe, 2002). What makes the mechanism proposed above so appealing is that it applies across multiple cellular processes; not just transcription. During its lifecycle, the virus must also utilize other cellular factors such as the DNA replication, repair, and recombination machinery (Dembowski and DeLuca, 2015; Muylaert and Elias, 2007; Taylor and Knipe, 2004). By utilizing nonspecific binding events as a means of attracting DNA-binding proteins and their cofactors, the virus has shifted the equilibrium in those locations, thereby enhancing the probability of regulatory factors binding at specific, functional sites. As a consequence, assembly of typically inefficient multi-protein complexes like the transcription pre-initiation complex (Darzacq et al., 2007), could become more favorable inside of RCs. We speculate that nonspecific protein-DNA interactions could be a general mechanism used by many other viruses. We also note that many RNA-binding proteins have been reported to undergo apparent LLPS (Courchaine et al., 2016) and believe it will be interesting to explore if RNA-binding proteins share a similar mechanism to what we describe here.

### Nonspecific binding events represent an important part of nuclear exploration

Our data also reveal a previously underappreciated aspect of how a DNA binding protein finds its target DNA within the nucleus. It has long been recognized that nonspecific binding to DNA could greatly accelerate the target search process by allowing for sliding in 1D along the DNA, thereby reducing the search space and allowing for faster-than-diffusion association kinetics (Berg et al., 1981). This is the case in bacterial systems where DNA is generally accessible to binding. A number of theoretical studies have addressed various aspects of the problem in eukaryotic systems, where TFs compete with nucleosomes for access to DNA (Mirny, 2010). In these cases, the DNA polymer is treated as a surface of variable binding energies on which a transcription factor may slide—if there are no nucleosomes to impede its diffusion (Mirny et al., 2009). *In vivo* experiments using sequence-specific eukaryotic transcription factors find that, for the cases studied, a given factor will spend approximately half its search time undergoing 3D diffusion, and the other half bound nonspecifically, presumably scanning in 1D (Chen et al., 2014; Hansen et al., 2017; Normanno et al., 2015).

The data we present here offer a new perspective on the importance of nonspecific and low-affinity binding, and competition for nucleosomes inside the nucleus. When the virus begins replicating, we measured the newly synthesized DNA to be ~130 times more accessible to DNA-binding proteins than the surrounding host chromatin (Table S3). Not only is there simply more DNA available to bind in the absence of nucleosomes, but the distance that a protein can scan in 1D is greatly increased due to the lack of impeding nucleosomes. Pol II is not a canonical DNA-binding protein, and no systematic study has been undertaken to measure its binding affinities against with different substrates. Still, having evolved to be an enzyme that must transcribe highly diverse DNA sequences suggests that its affinity for DNA outside of an assembled PIC may be relatively high. A recent study which explicitly modeled nonspecific DNA binding in the context of 3D genome organization finds that the most effective regime for recruiting a DNA binding protein is exactly what the virus appears to have settled on—that is a region of low overall DNA density that is free of nucleosomes recruiting a protein with high nonspecific affinity (Cortini and Filion, 2018).

The virus’ strategy may be shared by the host in facilitating enhancer-promoter contacts. Both promoter and enhancer elements are identifiable by their increased DNA accessibility. While some recent reports have suggested that enhancers may phase separate as a mechanism of activating transcription (Cho et al., 2018; Hnisz et al., 2017; Sabari et al., 2018), the data presented above suggest that the mechanisms that keep a cluster of enhancers and promoters accessible to DNA-binding proteins may facilitate the accumulation of Pol II and other PIC components, without the need for invoking LLPS.

### Mechanism of Pol II recruitment may explain robust transcription of late genes

An unresolved question in the study of herpesviruses is how genes with seemingly weak promoter elements can sustain such robust transcription, especially in the case of late gene promoters which often contain little more than a TATA box and Initiator elements (Rajčáni et al., 2004). How are these transcripts so highly expressed off of such weak promoters? While it is clear that other regulatory components also play a role in regulation of late genes (Davis et al., 2015; Lester and DeLuca, 2011; Li et al., 2018), our data may help shed light on how the virus robustly transcribes them after replication onset. After replication, when there are many copies of the viral genome present in a single RC, the compartmentalization of Pol II (and the other general transcription factors) mediated through nonspecific binding to the viral DNA also favors assembly of PICs at otherwise weak late gene promoters. In this way, the virus can conserve precious sequence space in its genome to encode other important features, and rely on fundamental mechanisms of nuclear exploration for Pol II, and other components of the transcription machinery, to provide sufficient gene expression for these late genes.

## Author Contributions

Conceptualization, D.T.M., X.D., and R.T.; Methodology D.T.M., A.S.H., H.M-N., S.T., X.D., and R.T.; Investigation, D.T.M., Y.H., K.K.U., and C.D-D; Software, D.T.M., A.S.H., H.M-N., S.T., and A.B.H; Writing – Original Draft, D.T.M; Writing – Review & Editing, D.T.M., A.S.H., H.M-N., S.T., A.B.H, C.D-D, X.D., and R.T.; Funding Acquisition, X.D., and R.T; Resources, C.D-D.; Supervision, D.T.M, X.D., and R.T.

## Acknowledgments

We would like to thank James Goodrich, Jennifer Kugel, and Robert Abrisch for providing the HSV1 strain KOS that began this project, and for helpful discussions. Thank you also to Neal DeLuca for the generous gift of the UL2/50 HSV1 strain. Thank you to Luke Lavis for generously providing all of the Janelia Fluor dyes that enabled these experiments. Thank you to Ana Robles and Astou Tangara for their tireless work keeping the microscopes in working order. Thank you to all of the individuals who provided reagents, comments, and critical insight for this manuscript, including Shasha Chong, Thomas Graham, Britt Glaunsinger, Ella Hartenian, Matthew Parker, and the Tjian and Darzacq Lab members. This work was supported by NIH grants UO1-EB021236 and U54-DK107980 (XD), the California Institute of Regenerative Medicine grant LA1-08013 (XD), by the Howard Hughes Medical Institute (003061, RT). A.B.H. is supported by the NIH predoctoral fellowship T32 GM098218. Portions of this work were performed on shared instrumentation at the CRL Molecular Imaging Center, supported by The Gordon and Betty Moore Foundation. We would like to thank Holly Aaron and Jen-Yi Lee for their assistance. DNA sequencing in this work used the Vincent J Coates Genomics Sequencing Laboratory at UC Berkeley, supported by NIH 669 S10 Instrumentation Grants S10RR029668 and S10RR027303.

## Supplementary Figure Legends

**Figure S1.**
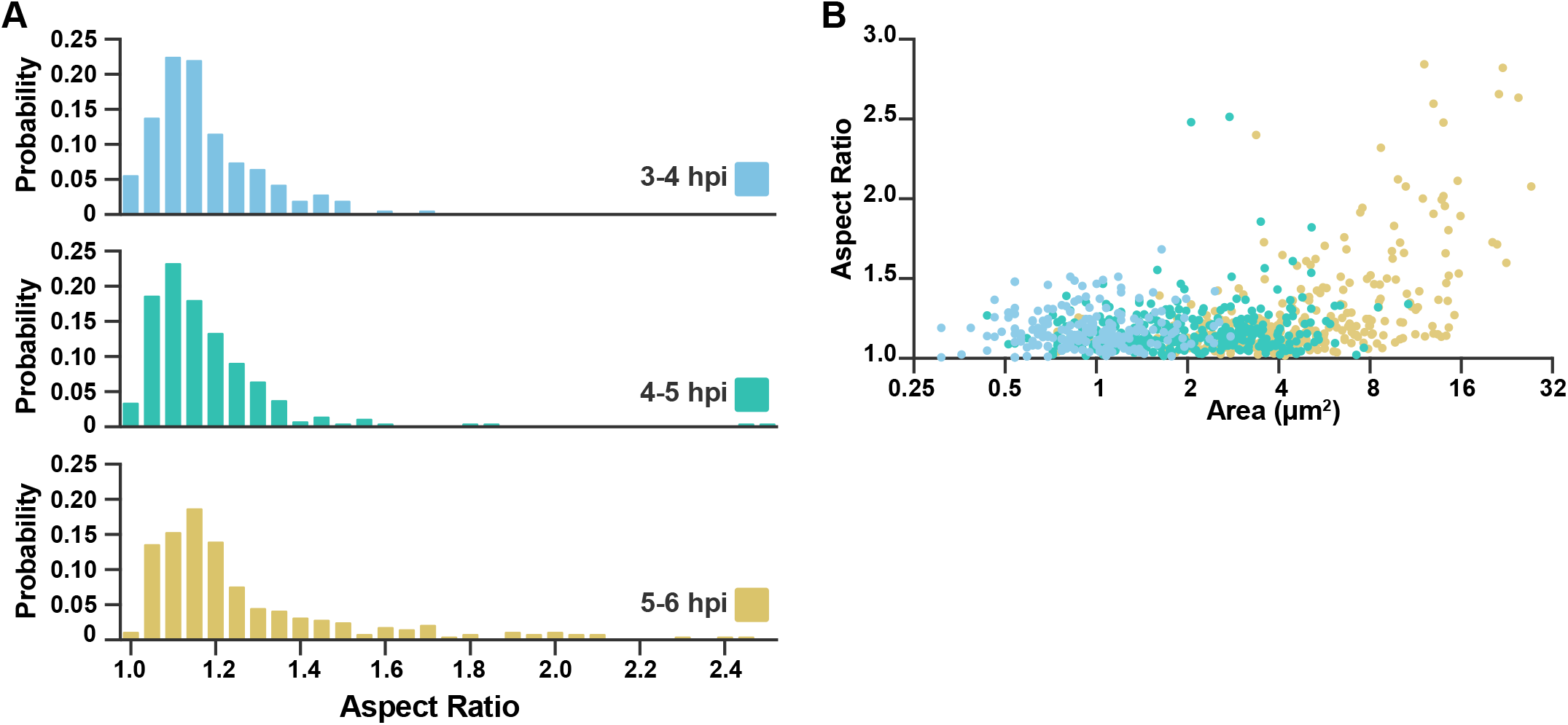
RCs remain round, particularly at early times in infection. Related to Figure 1. **A)** Histograms of the aspect ratios (Maximum diameter / minimum diameter) for RCs from cells at 3-4 hpi (n = 219), 4-5 hpi (n = 302), and 5-6 hpi (n = 296). **B)** Aspect ratio of every measured RC, as a function of its size measured in μm^2^. Dot colors correspond to the data sets in (A).

**Figure S2.**
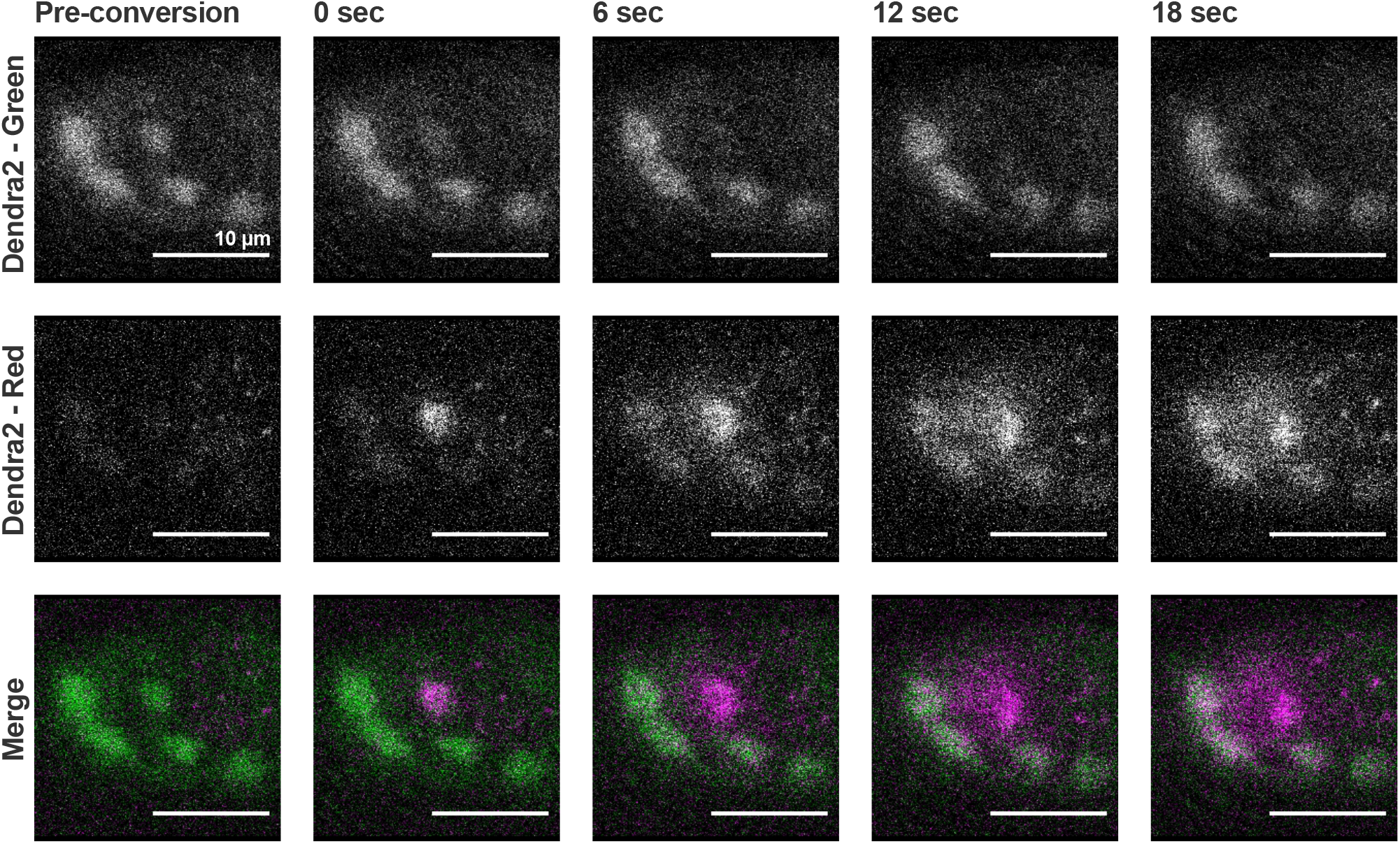
Dendra2 photoconversion shows Pol II exchanges with nucleoplasm. Related to Figure 2. Cells stable expressing Dendra2-Pol II were infected with HSV1. Fluorescence was monitored in both the green channel (pre-conversion), and red channel (post-conversion). A 1μm spot of 405nm light was used to convert one RC from green to red, alternating between photoconversion and frame acquisition. All scale bars are 10 μm.

**Figure S3.**
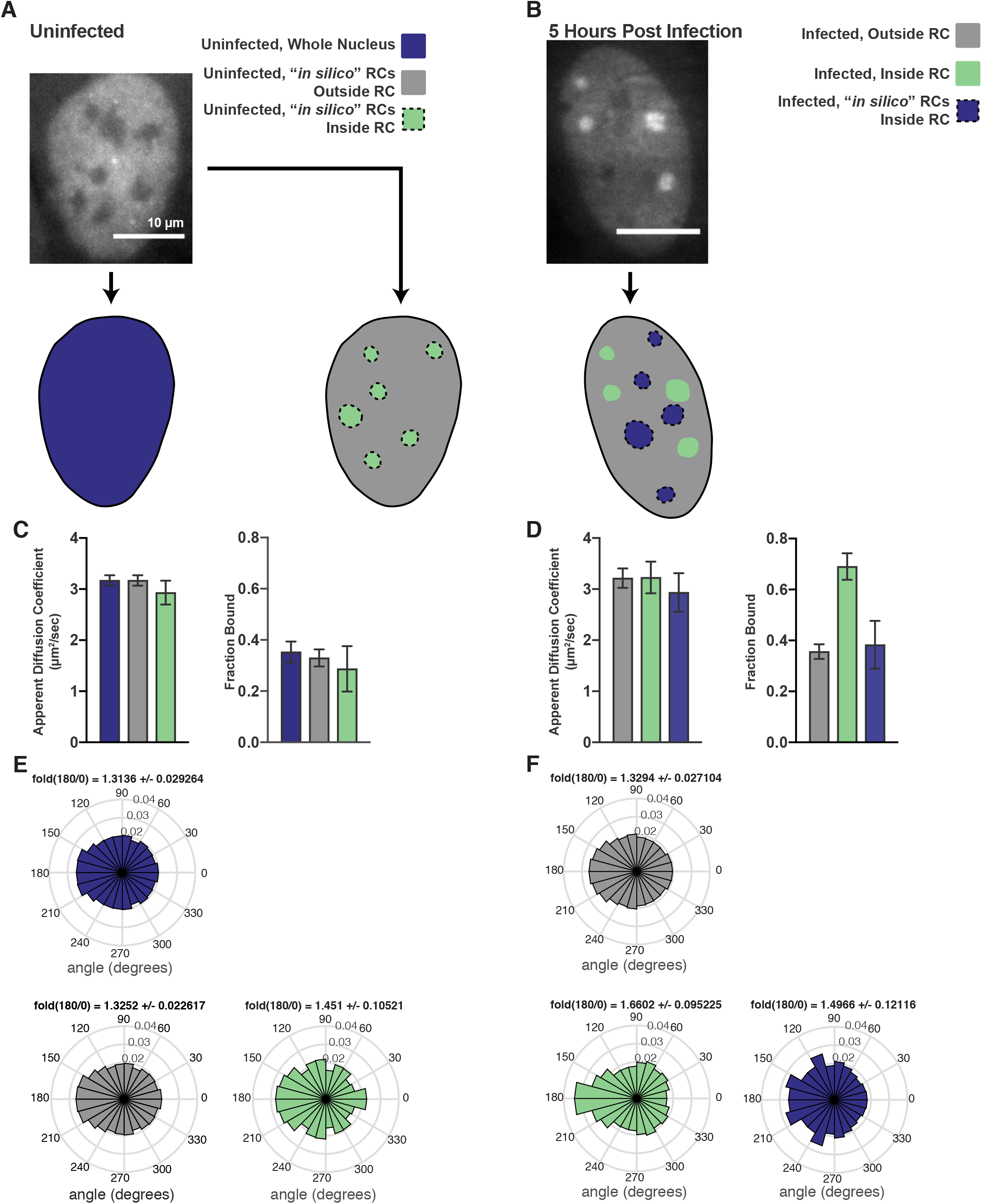
Comparison of *bona fide* RCs with RCs generated *in silico*. Related to Figure 2. **A)** Example workflow for uninfected cells, where either just the nucleus was masked (left), or the nucleus was masked and RC-sized annotations were randomly placed inside the nucleus (right). **B)** Example workflow for HSV1 infected cells, where both the correct annotations based on the widefield image and randomly shuffled RCs were generated for all measured cells. **C)** Spot-On measurements of trajectories after inside/outside classification in uninfected cells. *In silico* shuffling of RC positions has very little effect on either the measured apparent diffusion coefficient or the fraction bound. Error bars are the standard deviation of the mean, calculated from 100 iterations of randomly subsampling 15 cells without replacement and fitting with the model. **D)** Similar to (C), but for infected cells. Real RCs show an increase in fraction bound, whereas *in silico* shuffled compartments show no difference with trajectories outside RCs. **E)** Angular distributions of Pol II trajectories in the regions marked in (A) Fold(180/0) is the mean plus/minus the standard deviation, calculated from 100 iterations of randomly subsampling 15 cells without replacement and fitting with the model. **F)** Angular distributions of Poll II trajectories in the regions marked in (B). Fold(180/0) is the mean plus/minus the standard deviation, calculated from 100 iterations of randomly subsampling 15 cells without replacement and fitting with the model. All scale bars are 10μm.

**Figure S4.**
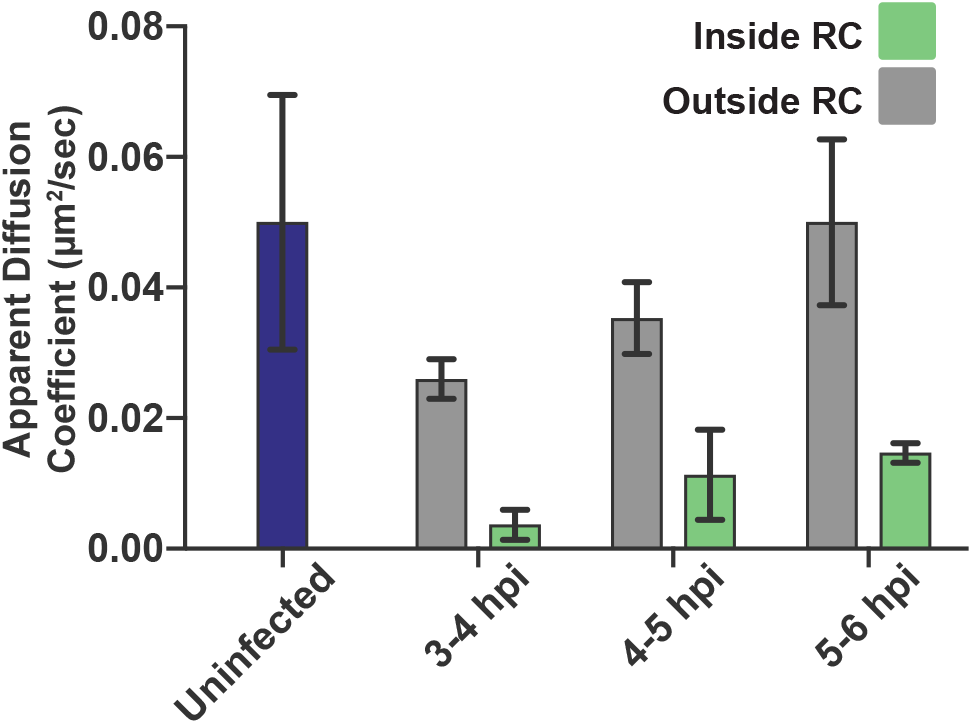
Diffusion coefficient of Bound population is consistent with chromatin binding. Related to Figure 2. Mean diffusion coefficient of the Bound population determined through 2-state model fitting for uninfected cells, and for cells at different times post infection, both inside and outside of RCs. In all data sets, the calculated diffusion coefficient is well below the upper bound set for the fitting, consistent with diffusion coefficients of chromatin (Hansen et al., 2018). Error bars are the standard deviation of the mean, calculated from 100 iterations of randomly subsampling 15 cells without replacement and fitting with the model.

**Figure S5.**
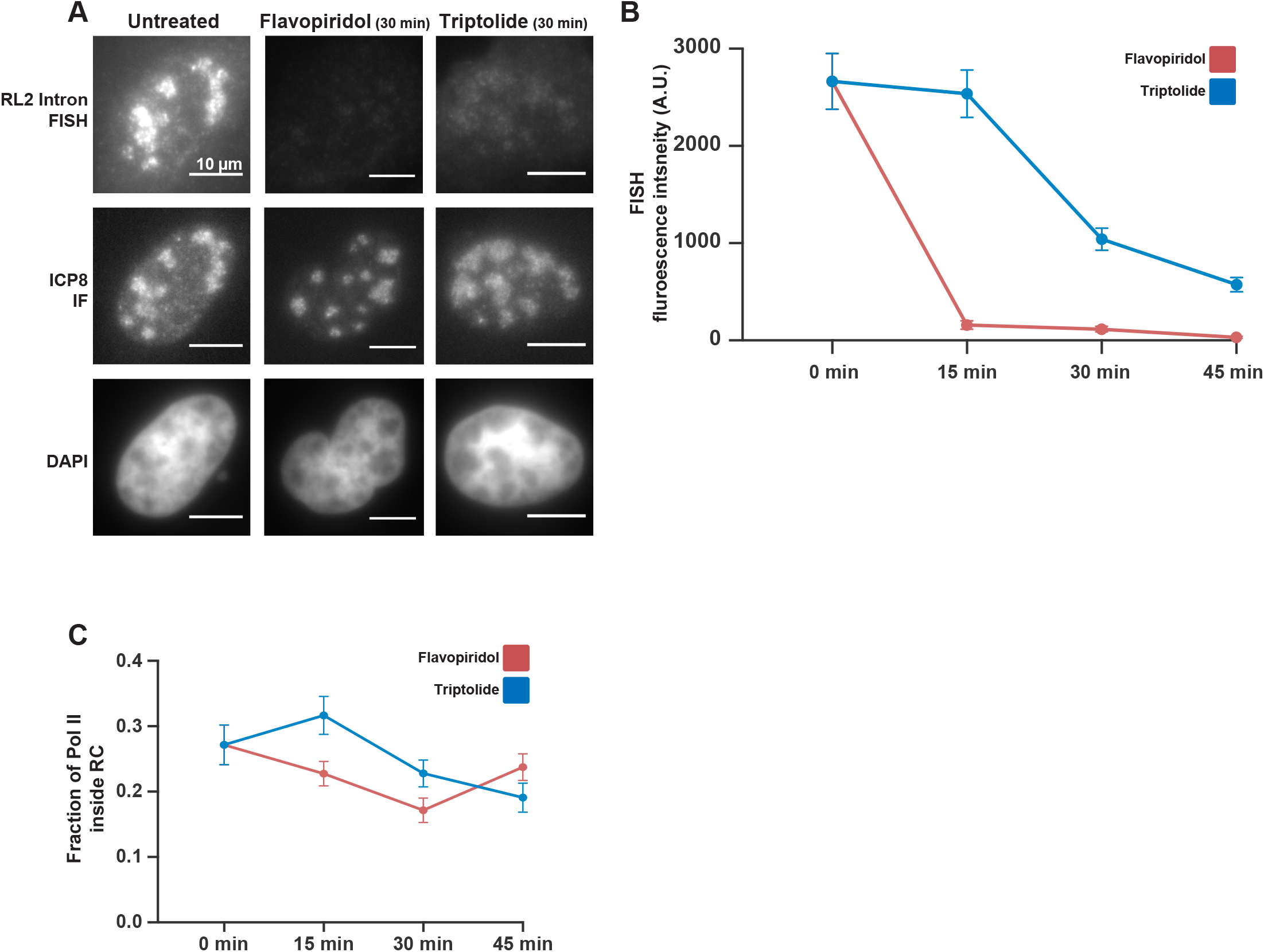
Transcription inhibition does not remove Pol II from RCs. Related to Figure 3. **A)** Representative images of U2OS cells, 4 hpi, either untreated or treated with the transcription inhibitors flavopiridol or triptolide. FISH probes against the intron to the HSV1 gene RL2 mark ongoing transcription, while immunofluorescence against the viral protein ICP8 marks infected cells by their RCs. All images within a color channel have been matched to have the same minimum and maximum pixel values. All scale bars are 10μm. **B)** Quantitation of transcription inhibition from (A). The FISH intensity in a 1μm z-section is compared in RCs from untreated cells (n = 170 RCs) to those treated for 15, 30, or 45 minutes with TRP (n = 192, 171, 191 RCs, respectively) and FLV(n = 158, 238, 153 RCs, respectively). Error bars represent standard error of the mean. C) Quantitation of Pol II recruitment to RCs from conditions in (A). The Pol II fluorescence intensity in a 1μm z-section is compared in RCs from untreated cells (n = 247 RCs) to those treated for 15, 30, or 45 minutes with TRP (n = 276, 206, 270 RCs, respectively) and FLV(n = 292, 246, 325 RCs, respectively). Error bars represent standard error of the mean.

**Figure S6.**
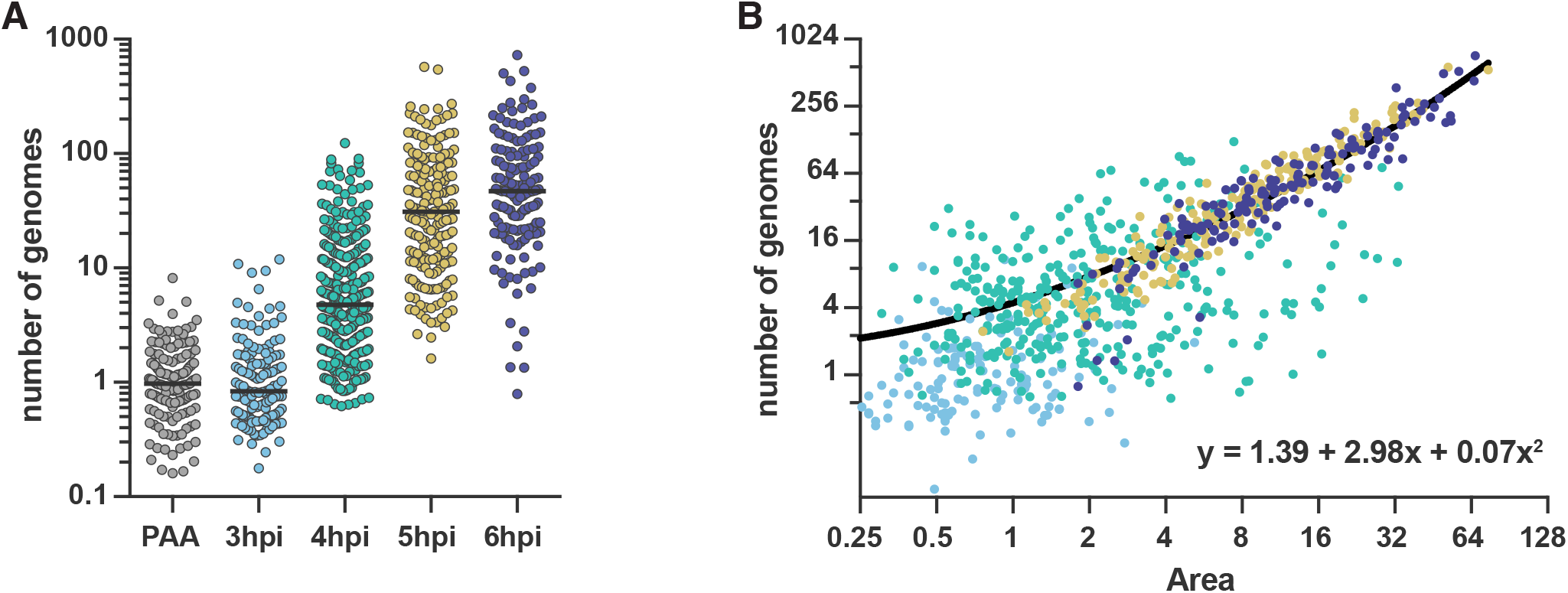
Quantification of DNA content in HSV1 RCs. Related to Figure 4. **A)** Dot plot showing the individual RC values from Figure 4C; normalized to the median of cells infected in the presence of PAA. Medians are indicated by a solid line (PAA = 1, 3 hpi = 0.83, 4 hpi = 4.77, 5 hpi = 31.13, 6 hpi = 46.95). **B)** Genome content of individual RCs in (A), plotted as a function of their area. Spot color corresponds to the times post infection as in (A). Line is a lease squares quadratic fit to all data points (R^2^ = 0.83).

**Figure S7.**
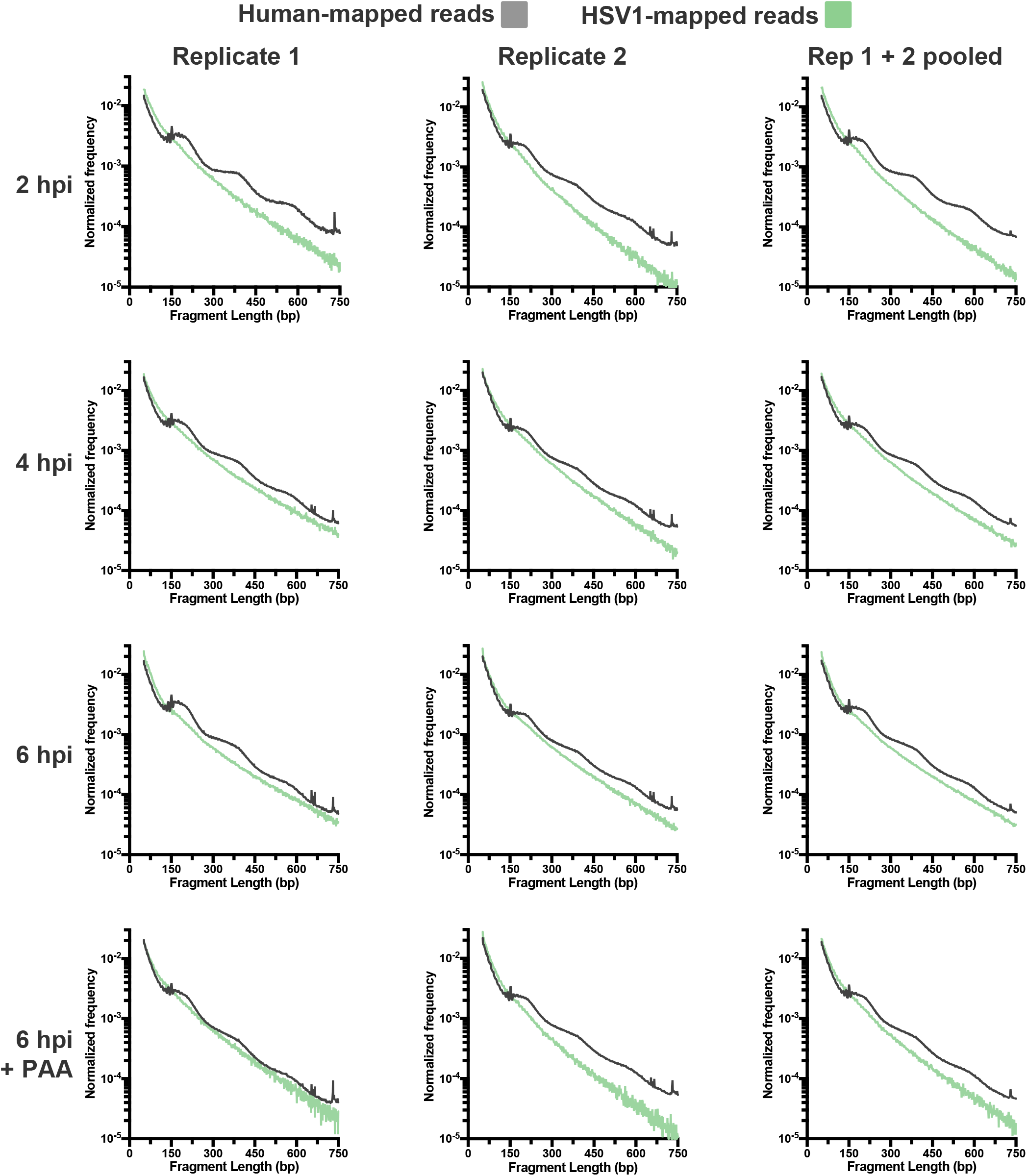
HSV1 genome appears nucleosome free at all times tested. Fragment length distributions of all conditions tested after HSV1 infection, for two individual replicates as well as for the pooled data. The green line indicates the lengths of fragments mapping to the viral genome, the gray line indicates lengths of fragments mapping to the human genome. All data are normalized to the total number of mapped reads to the respective genome, per condition.

**Figure S8.**
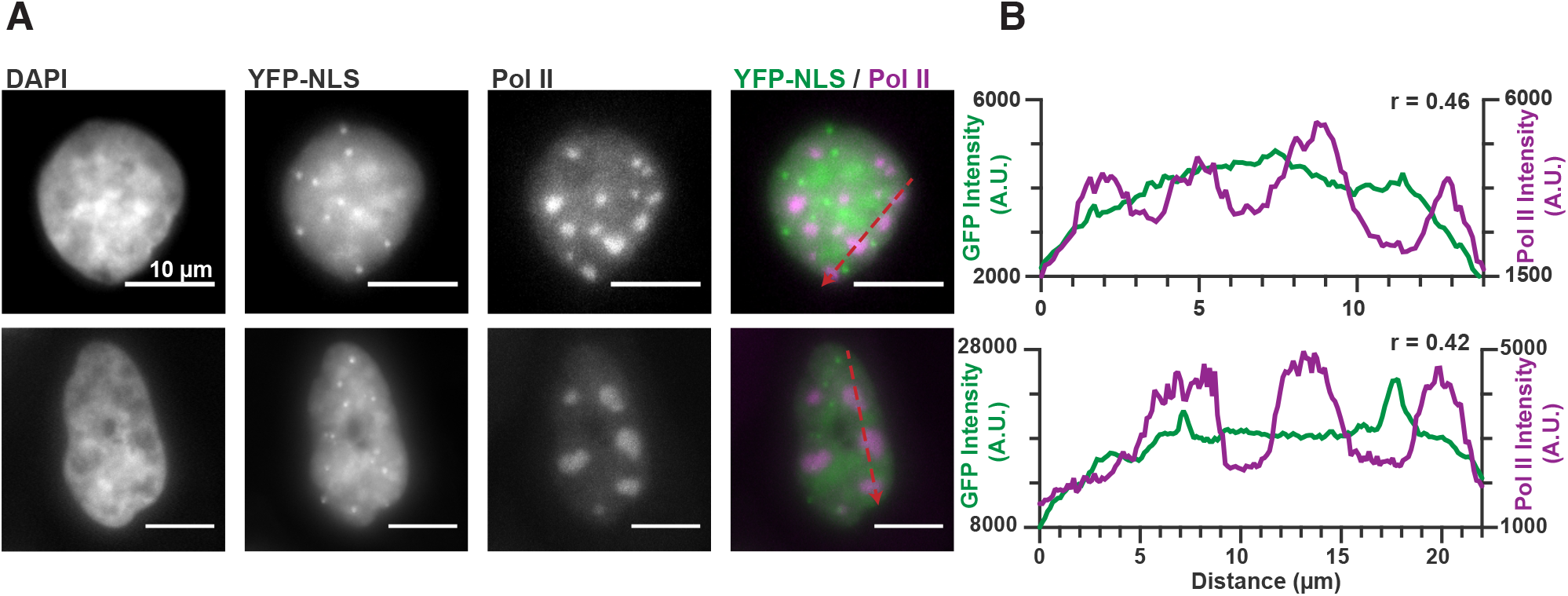
YFP-NLS does not colocalize with RCs. **A)** Representative images from cells transfected with YFP-NLS and then infected with HSV1. YFP-NLS forms occasional puncta in the nucleus, but these do not overlap with RCs, as marked by Pol II. **B)** Pixel line scans of images in (A). Red arrows in (A) give the direction of the x-axis. Left axis is the intensity of YFP-NLS fluorescence, right axis is the intensity of JF_549_-labeled HaloTag-Pol II fluorescence. Spearman’s correlation was used to measure the covariation in the two channels across the line. All scale bars are 10 μm.

**Table S1.**
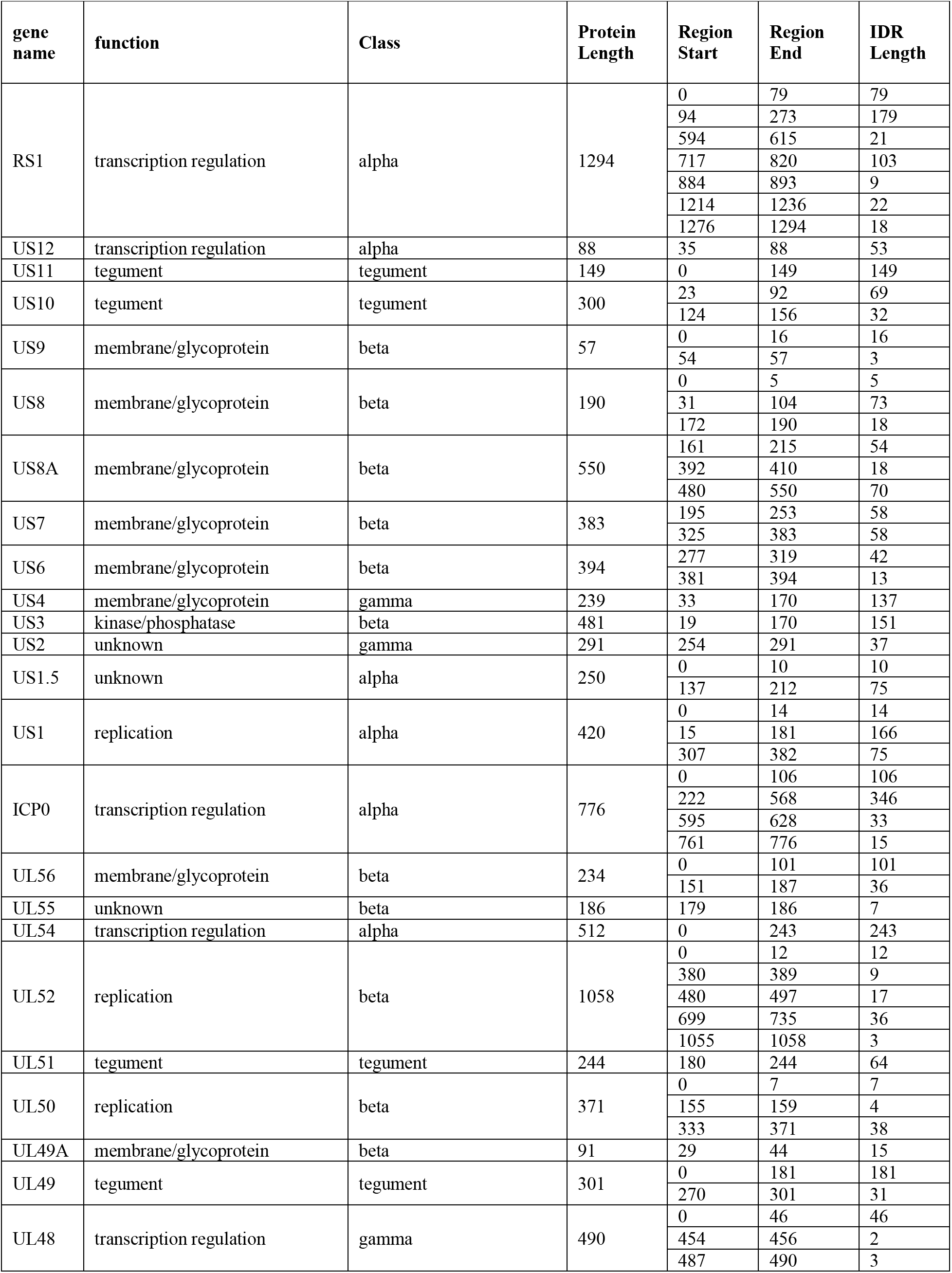
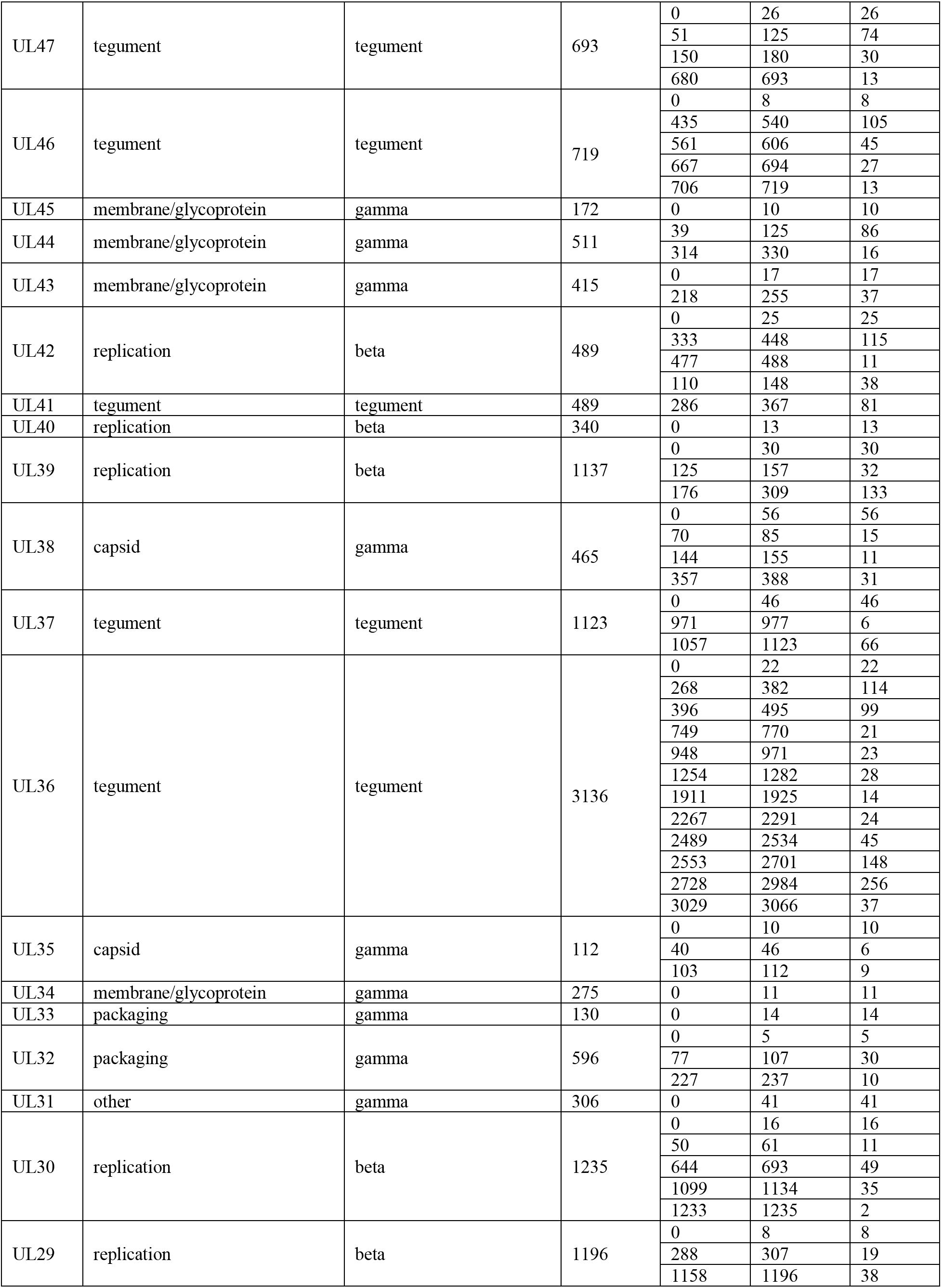
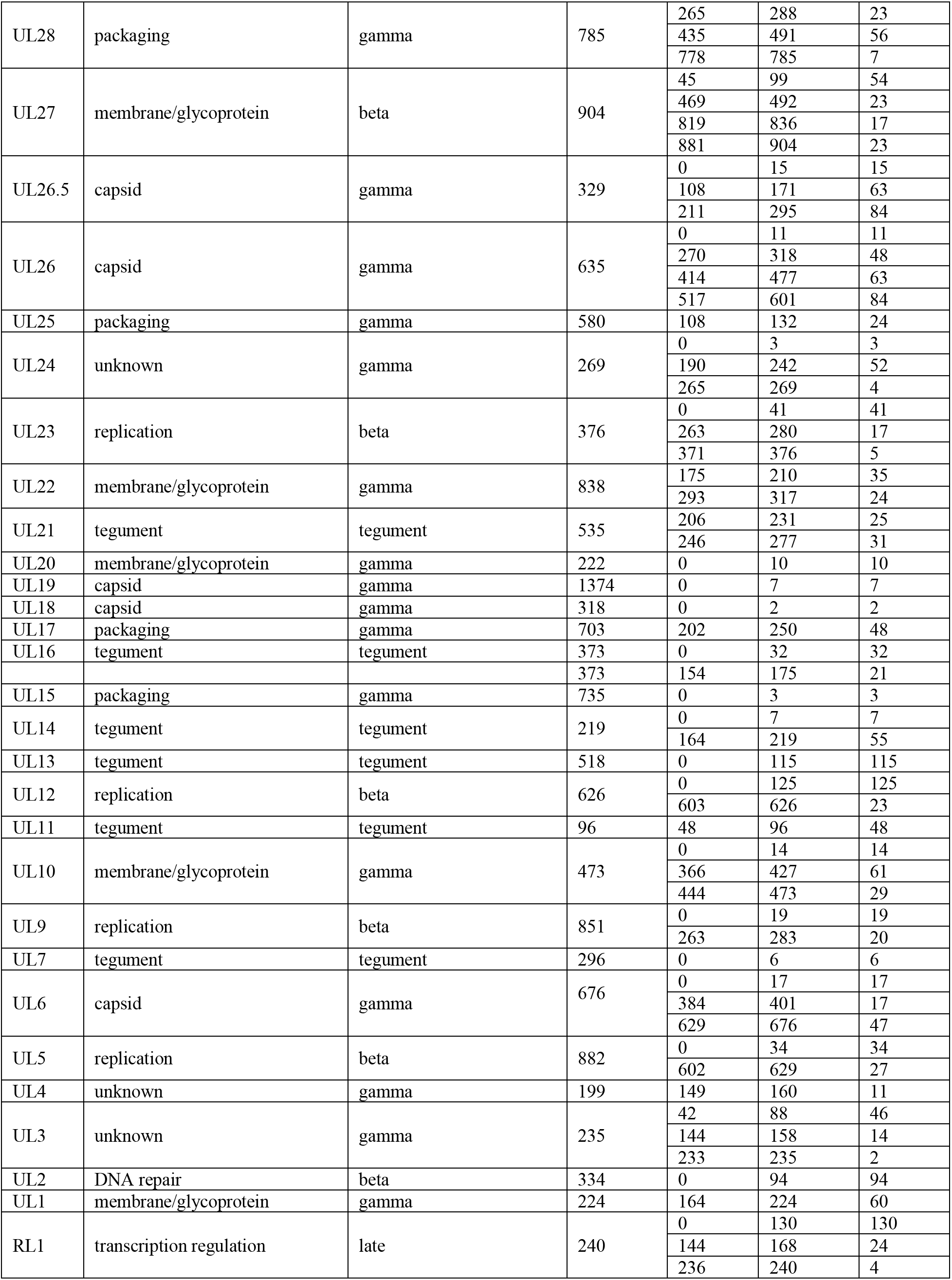
List of putative IDRs in the HSV1 genome identified by IUPred. Each protein listed was analyzed as described in the Methods section, and regions with an IUPred score of greater than 0.55 were recorded.

**Table S2.**
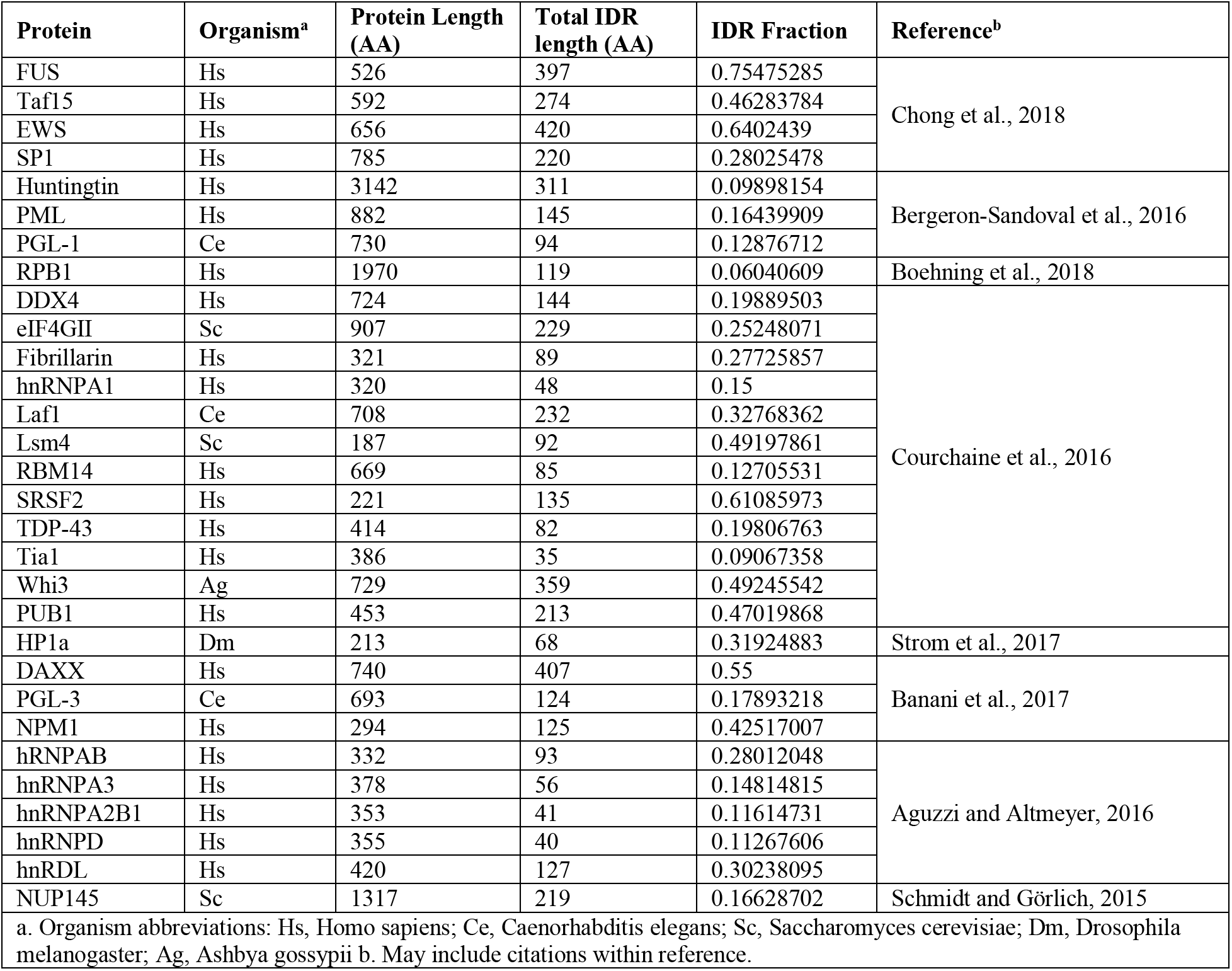
List of proteins reported to undergo phase separation. Gene name, organism of origin, size, and the fraction of the protein that scores as an IDR according to the analysis described in the Methods section. References and the citation within and provided.

**Table S3.** Quantitative measurements of HSV1 DNA inside of RCs. Using the values obtained through DNA FISH and ATAC-seq, we can make estimates of the copy number, concentrations, and relative enrichment of the viral DNA compared to the host. All values are calculated based on measurements of cells 6 hpi.

| **Table S3** | **Genome Size (bp)** | **Genome Copy number<sup>c</sup>** | **Total DNA (bp)** | **Percent of Total DNA<sup>c</sup>** | **Concentration (bp/$\mu\text{m}^3$)<sup>d</sup>** | **ATAC-seq read percentage<sup>e</sup>** | **Fold enrichment over expected<sup>f</sup>** |
| --- | --- | --- | --- | --- | --- | --- | --- |
| **Host Genome<sup>a</sup>** | 3.2 x10<sup>9</sup> | 2 | 6.4 x10<sup>9</sup> | 99.8 (± 0.2) | 9.4 (± 1.6) x10<sup>6</sup> | 75.8 (± 10.4) | 0.8 (± 0.1) |
| **Viral DNA** | 1.5 x10<sup>5</sup> | 82 (± 105) | 1.3 (± 1.6) x10<sup>7</sup> | 0.2 (± 0.2) | 3.9 (± 5.8) x10<sup>4</sup> | 24.2 (± 10.4) | 130 (± 170) |
| **Rel. Diff.<sup>b</sup>** | 2.1 x10<sup>4</sup> | | 513 (± 658) | | 240 (± 369) | | |

All values are the Mean ( ± S.D.).
a. Assuming karyotypically normal human cell; b. relative difference = Human / HSV1; c. Under experimental conditions of MOI = 1; d. Concentration assuming nucleus volume taken from Monier et al., 2000; e. based on total reads mapped from each organism, n = 3; f. Fold enrichment = ATAC-seq read percentage / Percent of Total DNA.

**Movie S1 and S2 – Time lapse movies of HaloTag-Pol II after HSV1 infection.** Cells were identified 3 hpi, and followed until they moved out of the focal plane.

## Experimental Procedures

### Tissue Culture

Human U2OS cells (female, 15 yr old, osteosarcoma) were cultured at 37°C and 5% CO2 in 1 g/L glucose DMEM supplemented with 10% Fetal Bovine Serum and 10 U/mL Penicillin-Streptomycin, and we subcultivated at a ratio of 1:3 – 1:6 every two to four days. Stable cell lines expressing the exogenous gene product α-amanitin resistant HaloTag-RPB1(N792D) or Dendra2-RPB1(N792D) were generated using Fugene 6 following the manufacturer’s protocol, and selection with 2 μg/mL α-amanitin. Stable colonies were pooled and maintained under selection with 1 μg/mL α-amanitin to ensure complete replacement of the endogenous RPB1 pool, as described previously (Boehning et al., 2018; Cisse et al., 2013).

Vero cells, were cultured for the growth and propagation of HSV1. Vero cells were cultured at 37°C and 5% CO2 in 4.5 g/L glucose DMEM supplemented with 10% Fetal Bovine Serum and 10 U/mL Penicillin-Streptomycin. Cells were subcultivated at a ratio of 1:3 – 1:8 every two to four days.

HSV1 Strain KOS was a generous gift from James Goodrich and Jennifer Kugel (Abrisch et al., 2015). UL2/50 was a generous gift from Neal DeLuca (Dembowski and DeLuca, 2015). All virus strains were propagated in Vero cells as previously described (Blaho et al., 2005). Briefly, cells were infected by incubation at an MOI ~ 0.01 in Medium 199 (Thermo) for one hour. 36-48 hpi, cells were harvested by freeze-thawing, pelleted, and sonicated briefly, and then centrifuged to clear large cellular debris. Because we were interested in the early events in infection, approximate titers were first determined by plaque formation assay in Vero cells (Blaho et al., 2005). More accurate MOI were determined by infecting U2OS cells plated on coverslips with the same protocol as would be using for imaging experiments. Cells were washed once with PBS, and then 100 μL of complete medium containing 1:10 – 1:10^5^ dilutions of harvested virus were added dropwise onto the coverslip to form a single meniscus on the coverslip. Infection was allowed to proceed for 15 minutes at 37 °C. Samples were then washed once with PBS and returned to culturing medium and incubated for 8 hours before fixation. To measure the MOI, immunofluorescence for the expression of ICP4 using an anti-ICP4 primary antibody (Abcam), and counting the number of infected versus uninfected cells. MOI was then calculated, assuming a Poisson distribution of infection events, as 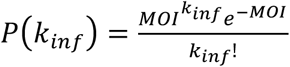, where kinf is the number of infection events per cell. When counting the uninfected cells, this simplifies to *MOI* = − ln(*f_uninfected_*). All experiments were performed from the same initial viral stock, with care taken so that each experiment was done with virus experiencing the same total number of freeze/thaw cycles to ensure as much consistency as possible.

### Live cell imaging

Cells were plated on plasma-cleaned 25 mm circular No. 1.5H cover glasses (Marienfeld High-Precision 0117650) and allowed to adhere overnight. For experiment using Halo-RPB1, cells were incubated with 50 – 500 nM fluorescent dye (e.g. JF_549_) conjugated with the HaloTag ligand for 15 minutes in complete medium. Cells were washed once with PBS, and the media replaced with imaging media (Fluorobrite media (Invitrogen) supplemented with 10% FBS and 10 U/mL Penicillin-Streptomycin). Prior to imaging, coverslips were mounted in an Attofluor Cell Chamber filled with 1 mL of imaging medium. Cells were maintained at 37 °C and 5 % CO2 for the duration of the experiment. For long term time course imaging experiments, cells were plated in 35mm No. 1.5 glass-bottomed imaging dishes (MatTek), infected with HSV1 at an MOI of ~1, and labeled with JF_549_, and finally the media exchanged for imaging media before placing in a pre-warmed Biostation (Nikon). At 3 hours post infection, infected cells were identified and imaged were taken every 30 seconds for 5 hours. For phase images, cells were plated and labeled as above, and imaged on a custom-built widefield microscope with a SLIM optics module (PhiOptics) placed in the light path directly before the camera.

### IUPred disorder prediction

Disorder predictions were preformed using a custom built python script to implement the IUPred intrinsic disorder prediction program (Dosztanyi et al., 2005; Dosztányi et al., 2005). Specific protein sequences were placed in a table and this was fed into the script. All protein sequences were downloaded from the reference organism at uniport.org. The resulting traces were smoothed by a rolling mean of 8 residues to remove noise and prevent single low-energy residues from splitting single large IDRs into multiple apparent IDRs. Contiguous substrings of residues with centered-mean IUPred disorder likelihood greater than 0.55 were annotated as “disordered regions” (Fig. 1E), and those contiguous regions larger than 10 amino acids were included in the calculation of “fraction IDR”.

### Fluorescence Recovery After Photobleaching (FRAP)

FRAP experiments were performed as previously described, with modifications. HaloTag-RPB1 cells labeled with 500 nM JF_549_ were imaged on an inverted Zeiss LSM 710 AxioObserver confocal microscope with an environment chamber to allow incubation at 37°C and 5% CO2. JF_549_ was excited with a 561 nm laser, and the microscope was controlled with Zeiss Zen software. Images were acquired with a 63x Oil immersion objective with a 3x optical zoom.

1200 total frames were acquired at a rate of 250 msec per frame (4 Hz). Between frames 15 and 16, an 11-pixel (0.956 μm) circle was bleached, either in the center of a RC, or in a region of the nucleus far from the nuclear periphery or nucleoli.

FRAP movies were analyzed as previously described (Hansen et al., 2017). Briefly, the center of the bleach spot was identified manually, and the nuclear periphery segmented using intensity thresholding that decays exponentially to account for photobleaching across the time of acquisition. We measured the intensity in the bleach spot using a circle with a 10 pixel diameter, to make the measurement more robust to cell movement. The normalized FRAP values were calculated by first internally normalizing the signal to the intensity of the whole nucleus to account for photobleaching, then normalizing to the mean value of the spot in the first 15 frames. We corrected for drift by manually updating a drift-correction vector with the stop drift every ~40 frames. FRAP values from individual cells were averaged across replicates to generate a mean recovery curve, and the error displayed is the standard error of the mean.

### Fluorescence Loss in Photobleaching (FLIP)

FLIP experiments were performed on the same microscope described above for FRAP.

Rather than bleach an 11-pixel spot a single time, in FLIP the spot is bleached with a 561 nm laser (or in the case of Dendra2, photoconverted with a 405 nm laser) between each acquisition frame. Movies were collected for 1000 frames at 250 msec per frame (4 Hz), or 1 frame per second (1 Hz) for Dendra2.

FLIP movies were analyzed using the same core Matlab code as the FRAP data, except that fluorescence intensities from another 10-pixel circle were recorded to measure the loss of fluorescence elsewhere in the nucleus. This analysis spot was chosen to be well away from the bleach spot, either at a neighboring RC in infected samples or somewhere else in the nucleoplasm far away from both the nuclear periphery and nucleoli. Instead of internally correcting for photobleaching, photobleaching correction was based on an exponential decay function empirically determined to be at a rate of e^−0.09^ per frame. FLIP data from multiple cells were averaged together to determine the mean and standard error for a given condition.

### RNA Fluorescence In Situ Hybridization (FISH) and immunofluorescence (IF)

RNA FISH was used to measure the transcription output for a given RC. To ensure we were measuring nascent transcription, we chose to tile the intronic region of RL2, one of the few HSV1 transcripts with an intron. The 25 oligonucleotide probes were synthesized conjugated with a Cal Fluor 610 dye (Biosearch Technologies). FISH was performed based on the manufacturer’s protocol. Briefly, cells were plated on 18 mm No. 1.5 coverslips (Marienfield) and infected. At the desired time point, cells were fixed in 4% Paraformaldehyde diluted in PBS for 10 minutes. After two washes with PBS, coverslips were covered with 70% v/v ethanol and incubated at −20 °C for 1 hour up to 1 week.

For hybridizations, coverslips were removed from ethanol and washed in freshly-prepared Wash Buffer A (2 volumes 5x Wash Buffer A, 1 volume formamide, 7 volumes H2O) (Bioseach Technologies). Hybridization buffer (10% v/v Dextran Sulfate, 300 mM Sodium Chloride, 30 mM Sodium Citrate, 400, 10% Formamide v/v, and 12.5 nM pooled fluorescent probes) was prepared freshly before each hybridization. A hybridization chamber was prepared with moistened paper towels laid in a 15cm tissue culture plate. A single sheet of Parafilm was laid over the moistened paper towel. 50 μL of hybridization buffer was pipetted onto the parafilm, and a coverslip inverted into the hybridization buffer. The chamber was sealed with parafilm and placed in a dry 37 °C oven for 4-16 hours. After hybridization, coverslips were placed back into a 12-well plate containing 1 mL Wash Buffer A and incubated twice for 20 minutes in a dry oven at 37 °C, with the second wash containing 300 nM DAPI. In a final wash step, cells were washed in Wash Buffer B (Biosearch Technologies). Coverslips were mounted on glass microscope slides in Vectashield mounting medium (Vector Laboratories) and the edges sealed with clear nail polish (Electron Microscopy Sciences). For experiments with combined immunofluorescence and FISH, primary antibody was added to the hybridization buffer at a concentration of 2 μg/mL. An additional wash step with Wash Buffer A containing 1 μg/mL antimouse polyclonal antibody conjugated to AlexaFluor 647 was performed before DAPI staining, and incubated at 37°C for 20 minutes.

Samples were imaged on a custom built epifluorescence Nikon Eclipse microscope equipped with piezoelectric stage control and EMCCD camera (Andor), as well as custom-built filter sets corresponding to the wavelength of dye used. All samples were imaged the same day after hybridaztion and/or incubation with secondary antibody, and all samples to be quantitatively compared across coverslips were imaged on the same day using exactly the same illumination and acquisition settings to minimize coverslip-to-coverslip variation.

### Single Particle Tracking (spaSPT)

Single particle tracking experiments were carried out as previously described (cit), but are described here in brief. After overnight growth, U2OS cells expressing Halo-RPB1 were labeled with 50 nM each of JF_549_ and PA-JF_646_. Single molecules imaging was performed on a custom-built Nikon Ti microscope fitted with a 100x/NA 1.49 oil-immersion TIRF objective, motorized mirror are to allow HiLo illumination of the sample, Perfect Focus System, and two aligned EM-CCD cameras. Samples were illuminated using 405-nm (140 mW, OBIS coherent), 561-nm (1 W, genesis coherent), and 633-nm (1 W, genesis coherent) lasers, which were focused onto the back pupil plane of the objective via fiber and multi-notch dichromatic mirror (405-nm/488-nm/561-nm/633-nm quad-band; Semrock, NF03-405/488/532/635E-25). Excitation intensity and pulse width were controlled through an acousto-optic transmission filter (AOTF nC-VIS-TN, AA Opto-Electronic) triggered using the camera’s TTL exposure output signal. Fluorescence emissions were filtered with a single bandpass filter in front of the camera (Semrock 676/37 nm bandpass filter). All of the components of the microscope, camera, and other hardware were controlled through NIS-Elements software (Nikon).

For all spaSPT experiments, frames were acquired at a rate of 7.5 msec per frame (7 msec integration time plus 0.447 msec dead time). In order to obtain both the population-level distribution of the molecules for masking and the single trajectories, we used the following illumination scheme: First 100 frames with 561 nm light and continuous illumination were collected; then 20,000 frames with 633 nm light at 1 msec pulses per frame and 0.4 msec pulses of 405 nm light during the camera dead time; then 100 frames with 561 nm light and continuous illumination were collected. 405 nm illumination was optimized to achieve a mean density of ~ 0.5 localizations per camera frame.

### spaSPT data processing

SPT data sets were processed in 4 general steps using a custom-written Matlab (Mathworks): 1) Masks for RCs were annotated manually, 2) the masks were corrected for drift throughout the sample acquisition, 3) particles were localized and trajectories constructed, and 4) trajectories were sorted as “inside” compartments or “outside”.

First, the 100 frames at the beginning and the end of each movie were separately extracted and a maximum-intensity projection used to generate “before” and “after” images of the cell or cells in the field of view. These images would be used to correct for movement of the cell as well as the individual RCs. For each cell, the nucleus was annotated in the “before” image, and then again in the “after” image. We assumed that the cell movement over the ~4 minutes of acquisition was approximately linear, and calculated the drift-corrected nuclear boundary for every frame in the stack of SPT images. The same procedure was applied to each of the replication compartments. Particle localization and tracking were implemented based on an adapted version of the Multiple Target Tracking (MTT) algorithm, available at https://gitlab.com/tjian-darzacq-lab/SPT_LocAndTrack. In the first step, particles were identified with the following input parameters: Window = 9 px; Error Rate = 10^−6.25^; Deflation Loops = 0. Following detection, a mask generated from the drift-corrected nuclear boundary was applied to discard any detections not within the nucleus. Trajectories were reconstructed with the following parameters: Dmax = 10 μm^2^/sec; Search exponent factor = 1.2; Max number of competitors = 3; Number of gaps allowed = 1.

Finally, after trajectories have been reconstructed, they were sorted as “inside” RCs or “outside”. To minimize the potential for bias in calling trajectories inside of compartments, we only required a single localization in a trajectory to fall within a compartment for that trajectory to be labeled as “inside”. As is discussed in the main text, we tested this sorting strategy for implicit bias by computationally generating mock RCs in uninfected or infected samples (Figure S3). To do this, all of the annotations for RCs from the infected samples (n = 817), as well as the distribution of number of RCs per infected cell, were saved in a separate library. We then took the uninfected cells and, in a similar process as described above, annotated the nuclear boundary and nucleoli. We then randomly sampled from distribution of RCs per cell a number of RCs to place in the nucleus, and then from the library of annotations randomly chose these RCs and placed them in the nucleus by trial-and-error until all of the chosen RCs could be placed in the nucleus without overlapping with each other, a nucleolus, or the nuclear boundary (Figure S3A). The SPT data were then analyzed as above—drift-correction, followed by localization, building of trajectories, and sorting into compartments—using the exact same parameters. We also followed this same procedure of randomly choosing and placing artificial RCs in infected cells, this time avoiding previously annotated RCs instead of nucleoli (Figure S3B).

### Two-state kinetic modeling using Spot-On

We employed the Matlab version of Spot-On (available at https://spoton.berkeley.edu) in our analysis, and embedded this code into a custom-written Matlab routine. All data for a given condition were merged, and histograms of displacements were generated for between 1 and 7 Δt. These histograms were fitted to a two-state kinetic model which assumes one immobile population and one freely diffusing population: Localization Error = 45 nm; Dfree = [0.1 μm^2^/sec, 5 μm^2^/sec]; Dbound = [0.001 μm^2^/sec, 0.3 μm^2^/sec]; Fraction Bound = [0, 1]; UseWeights = 1; UseAllTraj = 0; JumpsToConsider = 4; TimePoints = 7; dZ = 0.700. Trajectory CDF data were fit to a two-state model as first outlined by Mazza and colleagues, and expanded with implementation in Hansen and colleagues.

Because of the sparsity of the data we collected, we could not reliably generate single-cell statistics. In order to estimate the variability in the data, we implemented a random subsampling approach where 15 cells from a particular condition were randomly chose and analyzed. The Dfree, Dbound, and Fraction Bound were calculated for all trajectories, for trajectories inside of RCs, and for trajectories outside of RCs. This process was repeated 100 times, and the median values and standard deviations calculated and reported.

### Analysis of angular distribution

Angular distribution calculations were performed using a custom written routine in Matlab, implementing a previous version of this analysis (available at https://gitlab.com/anders.sejr.hansen/anisotropy). To analyze the angular distribution of trajectories in different conditions, we started with the list of trajectories generated above, annotated as either “inside” or “outside” of RCs. A trajectory of length N will have N-2 three-localization sets that form an angle, and so we built a matrix consisting of all consecutive three-localization sets. It is crucially important that only diffusing molecules be considered in the analysis, as localization error of bound molecules would skew all of the data to be highly anisotropic. To address this, we used two criteria. First, we only applied a Hidden-Markov Model based trajectory classification approach to classify trajectories as either diffusing or bound (Persson et al., 2013), and kept only the trajectories that were annotated as diffusing. Second, we applied a hard threshold that both translocations (1 to 2, 2 to 3) had to be a minimum of 150 nm, which ensured that we could accurately compute the angle between them. Because a particle may diffuse into or outside of the annotated region, we counted a trajectory as “inside” only if the vertex of the angle occurred within an annotated region.

### ATAC-seq

ATAC-seq experiments were performed as previously described (Buenrostro et al., 2013). Briefly, 100,000 U2OS cells stably expressing HaloTag-RPB1 were plated and allowed to grow overnight. The following day, cells were infected as described above, and incubated either in complete medium, or complete medium supplemented with 300 μg/mL phosphonoacetic acid (PAA). Infections were timed such that all cells were harvested at once. All of the infected cell lines were then trypsinized, and 100,000 cells were transferred to separate eppendorff tubes.

Cells were briefly centrifuged at 500 xg for 5 minutes at 4°C, and the supernatant discarded.

After one wash with ice-cold PBS and another 5 minute spin at 500 xg and 4°C, cells were resuspended directly in tagmentation buffer (25 μL 2x Buffer TD, 22.5 μL nuclease free water, 2.5μL Tn5 (Illumina)) and incubated for 30 minutes at 37 °C. DNA extraction and amplification with barcodes were performed as previously described, with 10-16 total cycles amplification. Barcoded samples were pooled in equimolar amounts and sequenced using a full flow-cell of an Illumina Hi-Seq 2500 per replicate. Two replicates were performed. Sequenced reads were mapped separately to hg19 genome using Bowtie2 (Langmead and Salzberg, 2012) with the following parameters: --no-unal --local --very-sensitive-local --no-discordant --no-mixed – contain --overlap --dovetail --phred33. Reads were separately mapped to the HSV1 genome, JQ673480, using Bowtie2 with the following parameters: --no-unal --no-discordant --no-mixed --contain --overlap --dovetail --phred33. The bam files were converted to bigwig files and visualized using IGV (Robinson et al., 2011). TSS plots were generated using Deeptools suite (bamCoverage, computeMatrix, plotHeatmap tools) using UCSC TSS annotations for hg19 genome, and using a highly refined map of the gene starts in HSV1 kindly provided by Lars Dölken (University of Cambridge, to be published separately).

### Oligopaint on infected cells

For DNA FISH experiments, custom pools of fluorescently labeled DNA oligos were generated using previously published protocols (Beliveau et al., 2015; Boettiger et al., 2016). Briefly, oligo sequences tiling a 10,016 bp region in the Unique Long arm (JQ673480 position 56,985 to 66,999) and a 7703 bp region in the Unique Short arm (JQ673480 position 133,305 to 141,007) were manually curated using oligo BLAST (NCBI) against the HSV1 and human genomes with the following settings, following guidelines for Tm, GC-content, and length from previous Oligopaint protocols (Beliveau et al., 2012; Boettiger et al., 2016). Individual oligos were synthesized and pooled. PCR was used to introduce a common T7 promoter on the 3’ end of the final probe sequence, then the PCR products were gel purified before *in vitro* transcription to generate ssRNA complimentary to the hybridization sequence. Finally, the entire RNA pool was reverse transcribed in a single reaction using Maxima RT (Thermo) using either AlexaFluor-647 or AlexaFluor-555 5’-labeled oligos as the reverse transcription primer. After acid hydrolysis to remove the RNA, oligos were purified using high binding capacity oligo cleanup columns (Zymo) and resuspended in TE.

Cells were plated on 18 mm coverslips and infected as described above. Infected was allowed to progress for between 3 and 8 hours in the presence or absence of phosphonoacetic acid, then fixed with 4% paraformaldehyde for 15 minutes. Coverslips were washed twice with PBS, then incubated with 100mM Glycine in PBS for 10 minutes. Samples were permeabilized for 15 minutes with 0.5% Triton-X100 in PBS, then washed twice with PBS. After permeabilization, samples were treated with 100 mM HCl for 5 minutes, then washed twice with PBS. Prior to hybridization, samples were washed twice with 2X SSC (300 mM NaCl, 30 mM Sodium Citrate), and then incubated at 42 °C for 45 minutes in 2X SSC with 50% v/v Formamide. Coverslips were inverted onto a slide containing 25 μL hybridization buffer (300 mM NaCl, 30 mM Sodium Citrate, 20% w/v Dextran Sulfate, 50% v/v Formamide, and 75 pmol of fluorescently-labeled oligos) and sealed with rubber cement. Samples were denatured at 78 °C on an inverted heat block for 3 minutes, then incubated in a humidified chamber at 42°C for 16 hours. Samples were then removed from the glass slides and washed twice to 60 °C with prewarmed 2x SSC for 15 minutes, then washed twice with 0.4x SSC at room temperature for 15 minutes. Finally, coverslips were mounted on glass slides with Vectashield mounting medium.

DNA FISH samples were imaged on the same microscope as described above for immunofluorescence and RNA FISH. Z-stack images were collected from all the way below the focal plane to all the way above the focal plane, with a step size of 100 nm. All samples were imaged on the same day using the same illumination and acquisition settings to minimize coverslip to coverslip differences.

### Analysis of Immunofluorescence, RNA, and DNA FISH

All cells were analyzed using a custom built Matlab script. First, a single image for each color channel was generated by automatically identifying the focal plane of the stack, and then integrating the pixel intensity for all pixels 1 μm above and below the focal plane. Nuclei were automatically segmented, but replication compartments could not reliable by detected using simple thresholding, and so each was manually annotated. A region of the image was selected to represent the black background, and the mean pixel value of this region was subtracted from every pixel in the image. After segmentation, the pixel values for each nucleus were recorded, as well as every RC within a given nucleus, and these were used to measure the signal within the RC, as well as the fraction of signal within compared to the rest of the nucleus (immunofluorescence only).

### Quantification of DNA content within RCs

DNA FISH data were compared with ATAC-seq data for the 6 hpi timepoint. Despite the fact that U2OS are hypertriploid, we based all the calculations on the DNA content of a diploid cell. As such, the values presented here likely represent an upper bound on the relative concentrations of host and HSV1 gDNA for our experiments. Precise volume measurements for nuclei were based on data from Monier et al., 2000, volumetric measurements for RCs were taken directly from the annotations of the DNA FISH data. Measurement uncertainty was propagated following standard practices outlined in Taylor, 1997.

### PALM of Pol II in RCs

For PALM experiments to precisely localize Pol II molecules within RCs, cells were labeled with 500 nM PA-JF_549_, and then infected as described above. Cells were fixed in 4% Paraformaldehyde in PBS, washed twice with PBS. Fluorescent 100 nm and 200 nmTetraspek beads were mixed in a 9:1 ratio then diluted 1000-fold in PBS. 100 μL was added to each coverslip and allowed to settle for 5 minutes, followed by 5 minutes of washing while rocking.

Coverslips were mounted in Attofluor Cell Chambers and covered with PALM imaging buffer (50 mM NaCl, 50 mM Tris pH 7.9, 2 mM Trolox) to reduce triplet-state blinking.

Samples were imaged on a custom-built Nikon Ti microscope equipped similarly to the microscope for single particle tracking, with some differences described here. An Adaptive Optics module (MicAO) and a removable cylindrical lens were placed in the light path ahead of the EM-CCD (Andor iXon Ultra 897) cameras in the left and right camera ports (respectively) of the microscope. Astigmatism for precise 3D localization was introduced using the Adaptive Optics system. The Adaptive Optics system was controlled through the MicAO software and calibrated on 200 nM Tetraspek beads based on the total photon yield and point spread function shape after iterative tuning of the deformable mirror. After optimization, a slight astigmatism in the vertical Zernike mode (Astigmatism 90° = 0.060) was added, and several z-stacks of 100 nM Tetraspek beads with 10 nm between slices to calibrate the PSF shape with the Z-position. 30,000 frames were acquired with the 561 nm laser line and increasing amounts of 405 nm illumination in order to keep the number of single molecules consistent across the duration of acquisition.

Spatial statistics were collected on cells using previously published methods (Boehning et al., 2018). First, cell boundaries and replication compartments were annotated as for spaSPT experiments (above). Particularly for small objects like RCs, edge correction is crucial for accurate spatial point pattern statistics. Given a set of detections P, we used the estimator *f* to correct for biases generated by points near the RC boundary:

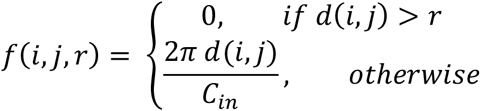

where *d*(*i,j*) is the distance between points *i* and *j* for *i,j*∈*P*, and C_in_ is arclength of the part of the circle of *d*(*i,j*) centered on *i* which is inside the annotated region (Goreaud and Pélissier, 1999). We then calculated N(r), the local neighborhood density:

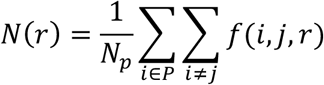

where N_p_ is the total number of detections within the region (Goreaud and Pélissier, 1999). The modified L-function is compared to complete spatial randomness (CSR), a homogenous Poisson process with intensity *λ*, equal to the density of detections in the region of interest A. The K-Ripley function is defined as:

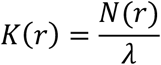

(Ripley, 1977). We estimated the modified L-function given by:

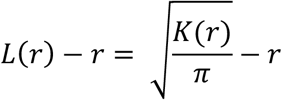

(Goreaud and Pélissier, 1999). For the modified L-function, a spatial distribution with CSR remains at 0 for all radii. To implement this analysis, we used a previously published python script and the ADS R package to estimate the spatial statistics (Boehning et al., 2018; Pélissier and Goreaud, 2015). In order to estimate the error in our measurements, for each cell we performed random subsampling of the data, before annotation, to randomly select 25,000 detections 100 times, and fed these subsampled data to the R script computing the statistic.

### STORM on infected cells

For STORM experiments to visualize both RNA Polymerase II and the viral DNA, U2OS cells stably expressing Halo-RPB1 were plated on coverslips, labeled with 300 nM JF_549_, and infected with the UL2/50 virus strain (Dembowski and DeLuca, 2015) as described above. After infection incubation with virus, cells were transferred into complete medium containing 300 μg/mL PAA for two hours to prevent replication. After two hours, cells were released from inhibition by exchanging the culture medium with complete medium containing 2.5 μM 5-Ethynyldeoxyuridine for 4 hours. Cells were fixed with 4% Paraformaldehyde in PBS for 10 minutes, then permeabilized with 0.5% Triton X100 in PBS for 10 minutes. Copper(1)-catalyzed alkyne-azide cycloaddition was performed with the ClickIT imaging kit following the manufacturer’s protocol (Thermo). Coverslips were mounted in Attofluor Cell Chambers and covered with freshly-made STORM buffer (50 mM NaCl, 50 mM Tris pH 7.9, 10% D-glucose, 10 mM DTT, 700 μg/mL Glucose Oxidase (Sigma), and 4 μg/mL catalase). STORM experiments were performed on the same microscope described for PALM.

### Data and Software Availability

The GEO accession number for the ATAC-seq data is: <u>GSE117335</u>. The SPT trajectory data are available via Zenodo at DOI:10.5281/zenodo. 1313872. The software used to generate these data is available at https://gitlab.com/tjian-darzacq-lab.

